# A multi-omics map of diabetic nephropathy reveals c-Jun as a driver of tubular injury and metabolic stress

**DOI:** 10.64898/2026.06.09.731164

**Authors:** Qiwen Deng, Yu Liu, Nathan A. Bracey, Yi-Chuan Wang, Paul Alexander Wadsworth, Marjia Afrin, Matteo Santoro, Shih-Yu Chen, Joseph C. Wu, Vivek Charu, Gerlinde Wernig

## Abstract

Diabetic nephropathy (DN) is a major cause of end-stage renal disease, yet the molecular mechanisms driving tubular injury and fibrosis remain poorly defined. Here, we integrated single-cell multiplexed protein imaging, spatial transcriptomics, single-nucleus and single-cell RNA sequencing and chromatin accessibility profiling to comprehensively characterize human DN pathology. Our multi-modal analysis precisely maps kidney cell types and their spatial distributions, immune-fibrotic interactions, and key transcriptional regulators. We identified eight distinct cellular neighborhoods defining the immune-fibrotic microenvironment and uncovered molecular networks driving tubular injury and fibrosis. *JUN* (encoding c-Jun) emerged as a central regulator of transcriptional reprogramming during tubular injury and fibrogenic remodeling. In a diabetic mouse model, c-Jun is activated in injured proximal tubules. Using an inducible c-Jun mouse model, we demonstrated that tubular-specific c-Jun activation alone is sufficient to induce tubular injury, chronic inflammation, progressive fibrosis, and systemic metabolic alterations, including impaired glucose homeostasis. We also observed reduced expression of SLC4A4, a bicarbonate transporter essential for proximal tubular function, in injured tubules. Together, our findings establish a spatially resolved framework for understanding DN pathogenesis and identify c-Jun as a key mediator of tubular injury and fibrosis in diabetic kidney disease.

## 1 Introduction

The annual healthcare costs of diabetes in the U.S. exceed $400 billion^1^. Diabetic nephropathy (DN) is a major complication of diabetes and accounts for nearly 50% of all cases of end-stage renal disease (ESRD)^2^. While DN shares pathological features with other chronic kidney diseases (CKD), including declining glomerular filtration rate (GFR), inflammation, and fibrosis^3^, the molecular mechanisms driving disease progression remain poorly understood, particularly within the complex kidney microenvironment. Advances in single-cell sequencing techniques such as scRNA-seq, snRNA-seq, snATAC-seq and spatial transcriptomics (e.g., Visium, CosMx) now enable high-resolution mapping of cellular and molecular alterations in human kidneys^4–8^. Bulk and single-cell transcriptomic studies have identified cellular heterogeneity in DN^4,6,9^, yet they lack spatial resolution to delineate how cellular interactions within fibrotic regions contribute to disease progression. Beyond transcriptomics, multiplexed imaging approaches provide protein-level insights into cellular states and interactions, offering a more comprehensive view of fibrotic remodeling and immune cell dynamics within tissue microenvironments.

For years, DN was considered a glomerulocentric disease, characterized by mesangial expansion, glomerular basement membrane thickening, and podocyte injury^10^, with tubular injury and interstitial fibrosis viewed as secondary consequences. However, accumulating evidence suggests that tubular injury can precede or occur concurrently with glomerular damage^11–14^. For instance, tubular injury markers such as kidney injury molecule-1 (KIM-1) and neutrophil gelatinase-associated lipocalin (NGAL) are elevated early in diabetes, often before the onset of significant albuminuria or detectable glomerular abnormalities^15–17^. Proximal tubule (PT) cells, essential for glucose reabsorption, are particularly susceptible to hyperglycemia-induced metabolic stress, leading to early dysfunction with long-term pathological consequences^18,19^.

Selective PT injury has been shown to initiate maladaptive responses in surrounding microenvironments, contributing to glomerulosclerosis and proteinuria^20^, reinforcing its role as a primary driver of DN progression. This paradigm shift highlights the critical role of tubule-specific mechanisms in DN pathogenesis and underscores their potential as therapeutic targets.

The tubules contribute to whole-body metabolic homeostasis through bicarbonate reclamation, gluconeogenesis and glucose reabsorption^21–24^. Tubular injury disrupts these processes, leading to metabolic acidosis, oxidative stress, and impaired glucose handling, which collectively accelerate kidney function decline^25–27^. Persistent metabolic stress and fibrosis further exacerbate tubular dysfunction^28,29^, reinforcing a cycle of maladaptation and chronic injury. While tubular injury is increasingly recognized as an important contributor to DN progression, the transcriptional programs and signaling pathways linking tubular injury to fibrosis remain poorly understood. To address these gaps, we employed single-cell multiplexed protein imaging (CODEX) and spatial transcriptomics (Visium) in 6 healthy and 24 DN kidney samples and integrated these datasets with single-cell/nuclear RNA sequencing and single-nucleus ATAC-seq to generate a comprehensive molecular atlas of the DN microenvironment.

This multi-modal dataset enabled high-resolution mapping of kidney cell types and their spatial organization, revealing the complex immune-fibrotic microenvironment in diabetic nephropathy. We identified functionally distinct cellular neighborhoods (CNs) including an injury enriched epithelial neighborhood marked by epithelial identity loss and mesenchymal-like features. Using pseudotime trajectory analysis, we delineated a continuum of proximal tubule cell states from homeostasis to injury-associated phenotypes, with c-Jun emerging as a key regulator of transcriptional reprogramming during tubular injury. In a diabetic mouse model, c-Jun activation was observed in injured proximal tubules. To further define the functional role of c-Jun *in vivo*, we generated a tubule-specific inducible mouse model in which c-Jun activation was sufficient to induce tubular injury, chronic inflammation, and progressive fibrosis. c-Jun activation was also associated with metabolic alterations, including impaired glucose homeostasis and suppression of renal gluconeogenic programs. Together, these findings generate a spatially resolved atlas of DN pathology and establish c-Jun-mediated transcriptional reprogramming as a key driver of tubular injury and fibrosis.

## 2 Results

### 2.1 Single-cell spatial proteomics reveals immune and fibrotic changes in DN

Multiplexed imaging allows us to capture a multitude of single-cell parameters while preserving their precise spatial context. To uncover the functional proteins in DN tissue, we created a specialized formaldehyde-fixed paraffin-embedded (FFPE) tissue microarray (TMA) for DN and employed CODEX with a panel of 51 markers. This strategy recreated key cellular phenotypes and the immune-fibrotic microenvironment landscape, enabling simultaneous profiling of 66 tissue regions from 22 participants (three regions each) (Fig. 1a-c, Extended Data Fig. 1a, b).

**Figure 1.**
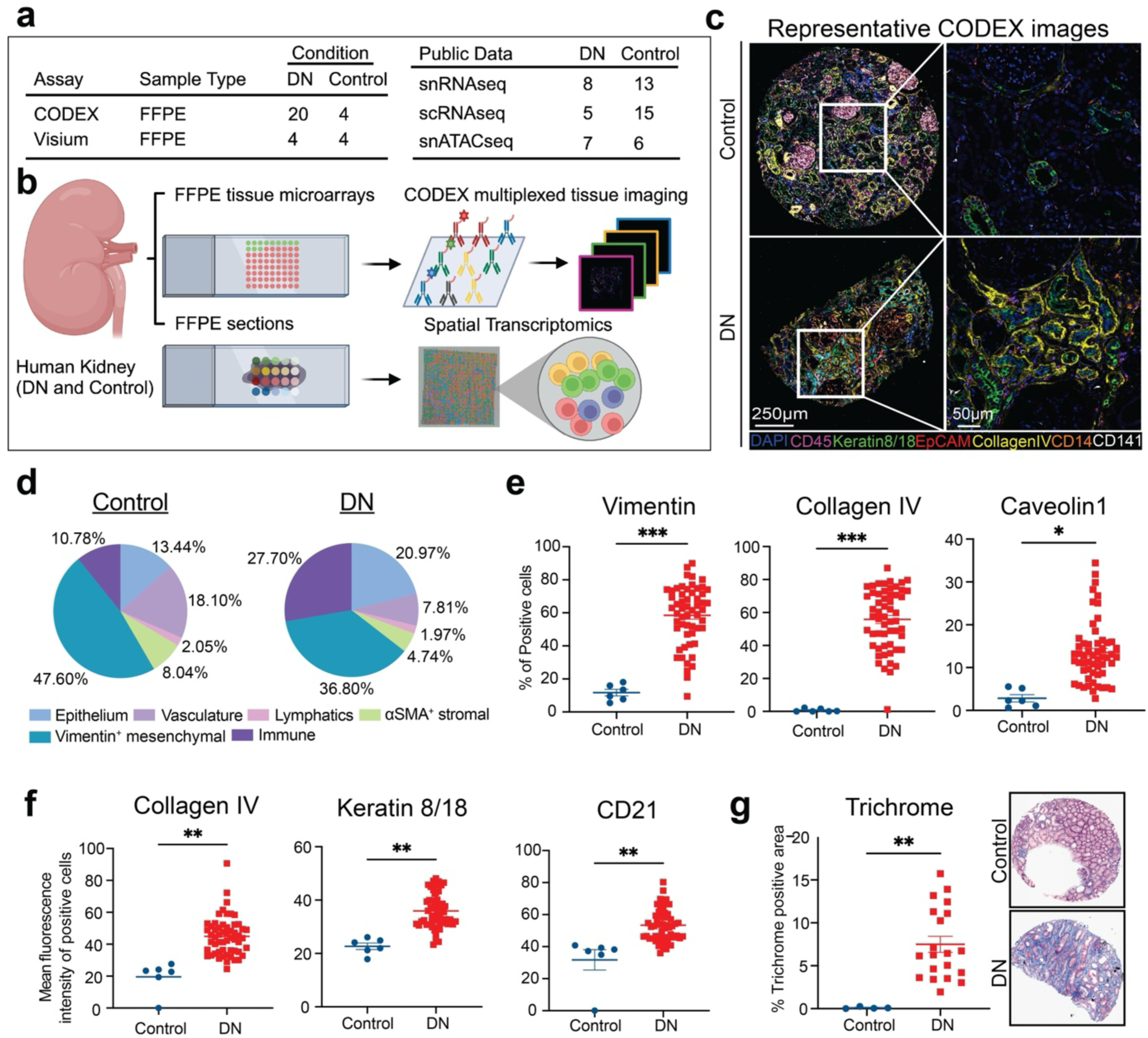
**Single cell multiplexed in situ imaging of human DN tissue**. a, Summary of the samples used in this study. DN, diabetic nephropathy. b, Tissues were processed for spatial transcriptomics (Visium) and CODEX multiplexed imaging assays. c, Representative CODEX images overlay (left) and overlay of representative seven markers (right). Scale bars, 250 µm (left) and 50 µm (right). d, Six main cell type clusters and their frequencies in all Control (n=12,174 cells) and DN (n=100,559 cells) samples calculated after CELESTA cell type assignment. e and f, Percentage of Vimentin, Collagen IV or Caveolin-positive cells and mean fluorescence intensity of Collagen IV, Keratin 8/18 or CD21-positive cells in glomeruli-excluded tissue regions from DN kidneys (56 regions across 20 samples) and Control kidneys (6 regions across 2 samples). For statistical analysis, the p-values were calculated based on the mean percentage per sample, with each sample containing between 1 and 3 regions. g, Trichrome staining of DN (n=20) and Control (n=4) samples.

Samples were split into two groups: healthy controls (n=2) including two peritumoral nephrectomies (without DN); and clinically diagnosed DN cases (n=20), with cohort characteristics summarized in Table 1 and individual clinical data detailed in Supplementary Table 1.

**Table 1.**
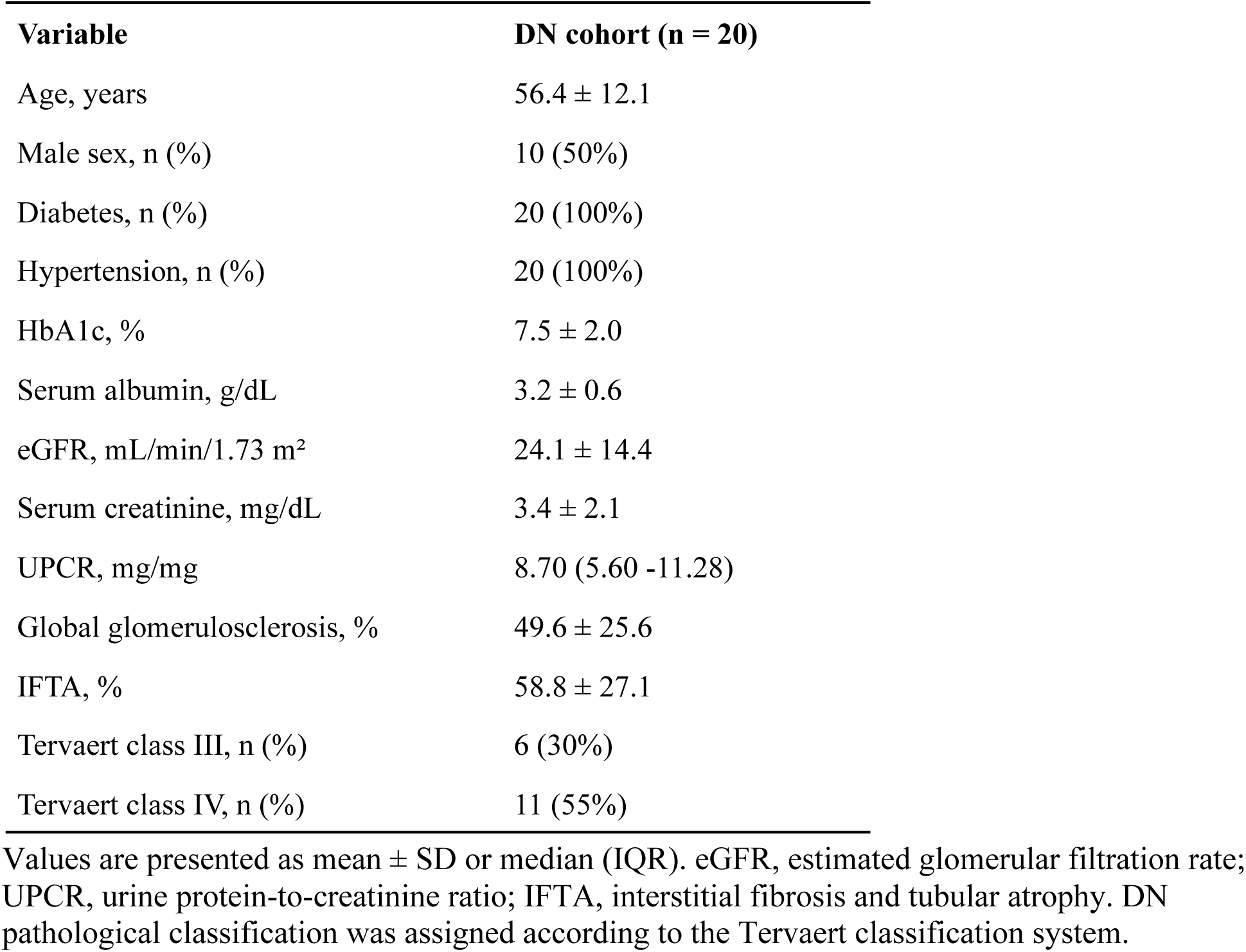
Clinical and pathological characteristics of the DN cohort used for CODEX analysis.

Using unsupervised clustering based on expression profiles and cell type assignment with CELESTA^30^, we distinguished six major cell types (Fig. 1d): epithelial cells marked by Keratin 8/18 and EpCAM; vascular cells defined by CD34 and CD31; lymphatics distinguished by Podoplanin; stromal cells highlighted by αSMA; mesenchymal cells identified by Vimentin; and immune cells characterized by CD45 (Extended Data Fig. 1a). Our protein maps revealed distinct differences between healthy and DN kidneys. DN tubules exhibited increased deposition of Vimentin, Collagen IV, and Caveolin 1 (Fig. 1e and 1f, *left*). Keratin 8/18, a well-established marker of tubular injury^31^, was upregulated in DN (Fig. 1f, *middle*). Additionally, CD21, typically a marker for B cells and follicular dendritic cells^32,33^, was detected in tubules and showed increased expression in DN samples (Fig. 1f, *right*). Trichrome staining further demonstrated a higher degree of fibrosis in DN tissues compared to healthy controls (Fig. 1g), highlighting widespread fibrogenesis and tubular injury. Furthermore, the co-expression of mesenchymal markers (Vimentin and αSMA) and epithelial markers (Keratin 8/18, E-cadherin, and EpCAM)—a signature of epithelial–to–mesenchymal transition (EMT), was far more pronounced in DN samples than in controls (Extended Data Fig. 1c). Lastly, quantification of immune cell clusters showed an increased presence of CD4^+^ T cells and CD68^+^ macrophages in DN, reflecting changes in the immune microenvironment (Extended Data Fig. 1d).

### 2.2 Characterization of the immune-fibrotic microenvironment (iFME) in DN

The iFME in DN represents a dynamic interplay between immune cells, resident kidney cells, and extracellular matrix (ECM) components that drive disease progression. By identifying spatial clusters, referred to as cellular neighborhoods (CNs), we analyzed the structural organization of this microenvironment. We qualitatively confirmed that cell type assignments aligned well with marker staining in the original CODEX images (Fig. 2a). To quantify colocalization, we calculated the average minimum distance (AMD) between cell types using SPIAT^34^. Since the measurement is not symmetrical, we specified the reference and target cell types used (Extended Data Fig. 2a). This analysis revealed several cell type pairs that were significantly more colocalized (Fig. 2b, *blue dots*) or more distant (Fig. 2b, *pink dots*) in DN compared to control kidneys. For instance, epithelial cells and CD68^+^ macrophages, as well as B cells with CD68^+^ macrophages and lymphatics, showcased striking differences. Representative CODEX images showed that Keratin 8/18^+^ epithelium shows increased spatial association with CD68^+^ macrophages (Fig. 2c), and CD20^+^ B cells display enhanced colocalization with CD68^+^ macrophages and Podoplanin^+^ lymphatic cells in DN compared to control samples (Extended Data Fig. 2b).

**Figure 2.**
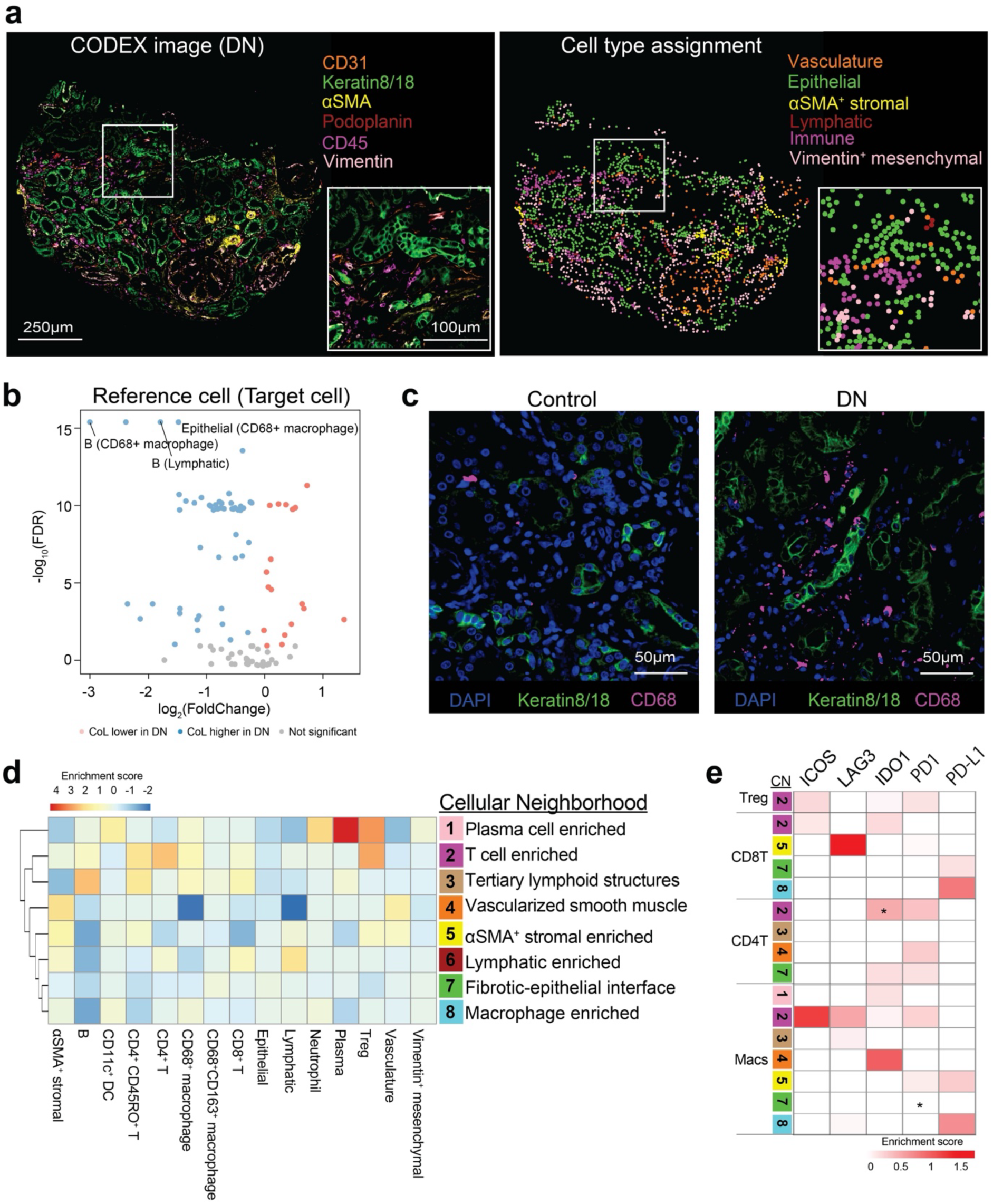
Decoding the immune fibrotic microenvironment in DN. a, CODEX image overlay (left) and CELESTA cell type assignment (right) for a representative TMA spot (donor no. 35379). Scale bars, main image, 250 µm; inset, 100 µm. b, Average minimum distance (AMD) between reference cells and target cells. CoL refers to how much two cell types are colocalized and potential interacting when comparing DN kidneys (56 regions across 20 samples) with Control kidneys (6 regions across 2 samples) in glomeruli-excluded tissue regions. c, Representative regions of a Control and DN sample shown as three-color overlay images. Scale bar, 50 µm. d, Identification of eight distinct cellular neighborhoods (CNs) based on 15 cell types and their respective percentage of positive cells in each CN. All samples are pooled together. e, The heatmap displays the mean enrichment of checkpoint-positive immune cell types, including T regulatory cells (Tregs), CD8^+^ T cells, CD4^+^ T cells and CD68^+^ macrophages, across different CNs. Asterisks denote significant differences in the proportions of checkpoint-positive cell types in specific CNs compared to the overall cellular proportions of the corresponding cell types across TMA cores (*p < 0.05, **p < 0.01, ***p < 0.001; Student’s t-test).

We applied a hierarchical clustering algorithm via SPIAT to identify CNs based on Euclidean distances between cells. Pairs of cells within a 50-unit radius were considered as interacting, while those beyond this threshold were considered non-interacting. Clustering was performed using data from both DN and control kidneys to identify common CNs. This analysis revealed eight distinct CNs that recapitulated the iFME of DN (Fig. 2d, Extended Data Fig. 2c). We validated CN assignments by overlaying them onto original CODEX images, confirming their alignment with marker staining (Extended Data Fig. 2d). The identified CNs reflected distinct tissue structures associated with disease progression. For example, we distinguished two stromal neighborhoods: CN4, enriched in vascularized smooth muscle, and CN5, marked by immune cell-infiltrated fibrotic stroma. CN6 is a hub of lymphatic cells, αSMA^+^ stromal cells, and both CD8^+^ and CD4^+^ T cells, reflecting lymphangiogenic events observed in human DN and other CKDs^35,36^. CN7 was enriched in epithelial cells, CD68⁺CD163⁺ macrophages, and Vimentin⁺ mesenchymal cells, consistent with a pro-fibrotic microenvironment characterized by active tubular injury, M2 macrophage-driven fibrogenesis, and excessive ECM deposition in renal fibrosis progression^37^. In addition, our analysis identified previously underestimated substructures in DN. For instance, CN3 was enriched in B cells, CD4⁺CD45RO⁺ T cells, CD8⁺ T cells, and CD68⁺ macrophages while being depleted of other cell types, resembling tertiary lymphoid structures (TLS). TLS are commonly found in IgA nephropathy^38^, lupus nephritis^39^, and transplanted kidneys^40,41^ but have been rarely described in DN. Other notable examples include CN1, enriched in plasma cells; CN2, enriched in T cells; and CN8, enriched in macrophages (Fig. 2d).

In addition to quantifying the proportion of each immune cell type within total immune cells, the analysis also allowed us to investigate their functional states by measuring the expression of five key immune checkpoint molecules—ICOS, LAG3, IDO1, PD1, and PD-L1 (Extended Data Fig. 2e). We observed highly expression of PD-1 and IDO1 on CD4⁺ and CD8⁺ T cells, PD-L1 on CD8^+^ T cells and CD68^+^ macrophages, and ICOS and LAG3 on CD4^+^ T cells. Given the distinct microenvironments within CNs, immune cells are expected to exhibit different functional states depending on their spatial context. We found that IDO1⁺ CD4⁺ T cells were significantly enriched in CN2, whereas PD-1⁺ CD68⁺ macrophages were significantly underrepresented in CN7, suggesting that diminished PD-1 expression may enhance their pro-fibrotic functions. This data highlighted the complex regulation of immune responses within the iFME of DN.

### 2.3 Spatial transcriptomics defines PT injury niches in DN

We employed the FFPE-Visium platform to unravel the molecular landscape and spatial organization of kidney cell types in DN. We generated eight spatial transcriptomics datasets, including four healthy (Control) and four disease (DN) samples (Supplementary Table 2). Given that each spatial transcriptomics spot captures a cluster of cells (∼55 μm in diameter), we computationally deconvoluted the cell-type composition of each spot using reference single-cell RNA-seq data (see Methods, Extended Data Fig. 3a–c). This enabled us to recapitulate cell types within each nephron segment aligning with histopathological features observed (Fig. 3a).

**Figure 3.**
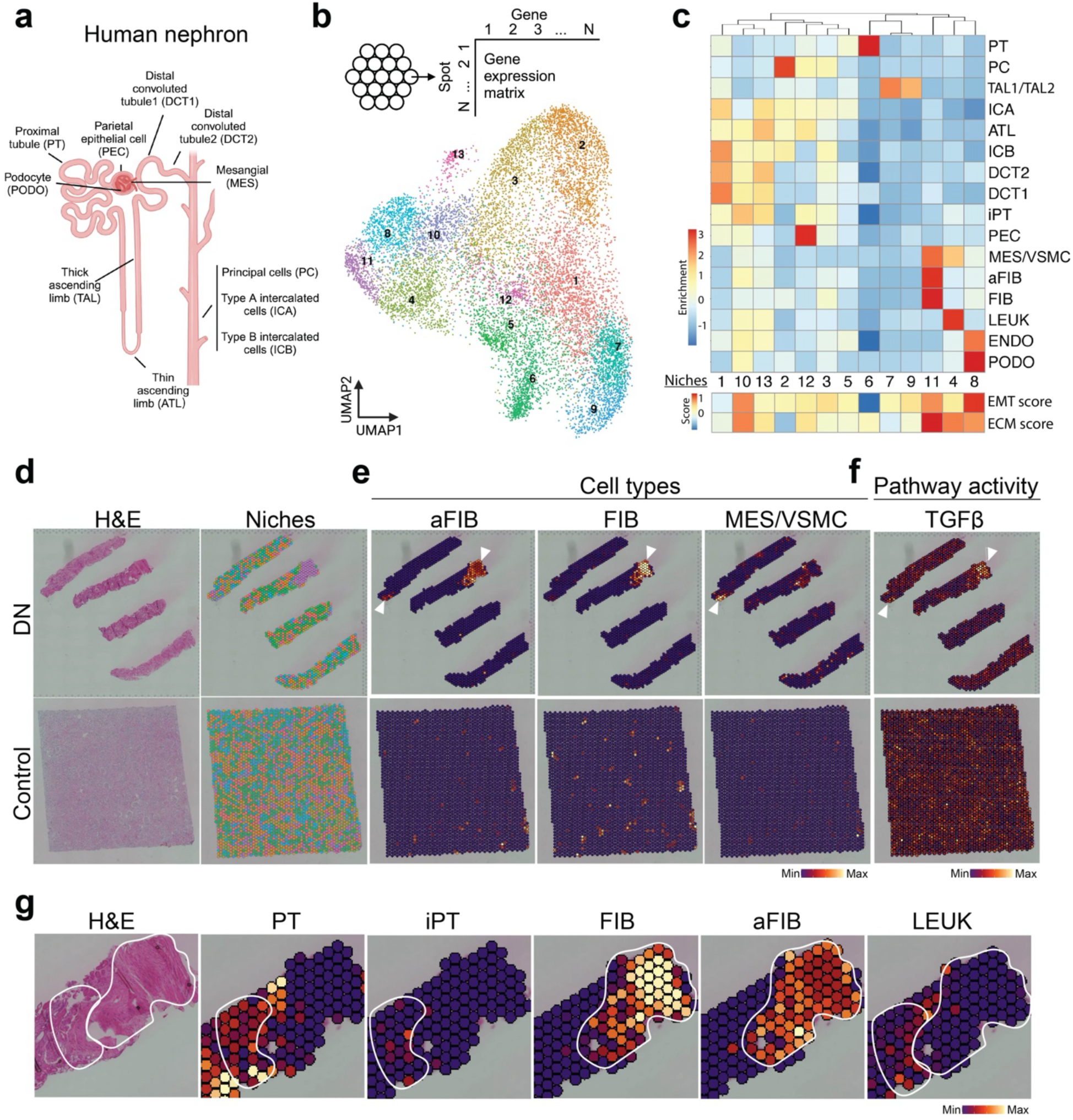
Spatial mapped molecular profile of human DN. a, Schematic of the human nephron showing cell types. b, Schematic of the cell niche and UMAP of spatial transcriptomics spots based on gene expression. c, Cell type compositions within each niche and corresponding EMT and ECM score for each niche. PT, proximal tubule; PC, principal cells; TAL1/TAL2, thick ascending limb; ICA/ICB, type A/B intercalated cells; ATL, thin ascending limb; DCT1/DCT2, distal convoluted tubule; iPT, injured proximal tubule; PEC, parietal epithelial cells; MES/VSMC, mesangial cells/Vascular smooth muscle cells; aFIB, activated fibroblasts; FIB, fibroblasts; LEUK, leukocytes; ENDO, endothelial cells; PODO, podocytes. d, H&E staining and spatial mapping of cell niches. e, Visualization of aFIB, FIB and MES/VSMC cell types and f, pathway activity for Control and DN tissues. g, Zoom-in image of H&E staining and spatial mapping of PT, iPT, FIB, aFIB and LEUK cells.

By clustering spots based on their gene expression profiles, we identified distinct molecular niches reflecting key cellular processes in DN (Fig. 3b, c, Extended Data Fig. 3d). Two glomerular niches (Niches 4 and 8) were associated with fibrotic and inflammatory process, enriched in endothelial cells (ENDO), podocytes (PODO), mesangial/vascular smooth muscle cells (MES/VSMC), and leukocytes (LEUK). A distinct fibrotic niche (Niche 11) contained a high proportion of fibroblasts (FIB), activated fibroblasts (aFIB), and MES/VSMC. We also identified three tubular niches: two injury-associated niches (Niches10 and 13) contained a high proportion of injured proximal tubule cells (iPT), distal convoluted tubule cells (DCT1/DCT2) and leukocytes, and one structural distal tubule niche (Niche 1) enriched with DCT cells and intercalated cells (ICA/ICB). Beyond these, we identified other structural niches, including a principal cell (PC) niche (Niche 2), a parietal epithelial cell (PEC) niche (Niche 12), a proximal tubule niche (Niche 6), and two thick ascending limb (TAL1/TAL2) niches (Niches 7 and 9). To further characterize these niches, we developed an ECM score based on the expression levels of collagens, glycoproteins, and proteoglycans^42^ and an EMT score incorporating EMT-related genes^43^. Notably, Niches 4, 8, and 11 exhibited high EMT and ECM scores, correlating with the enrichment of fibrotic cell types in these regions. Furthermore, Niche 10, predominantly composed of iPTs, also displayed elevated EMT and ECM scores, suggesting a direct link between proximal tubule injury and fibrotic remodeling (Fig. 3c).

The spatially resolved, cell-type-specific data allowed direct visualization of histological injury regions within patient biopsies (Fig. 3d). For example, regions enriched in aFIB, FIB, and MES/VSMC clearly marked fibrotic zones (Fig. 3e, *white arrow*). These fibrotic areas were adjacent to atrophic tubules (Fig. 3g, PT and iPT, *white circle)* and surrounded by immune cells (LEUK, *white circle*), which exhibited both injury and profibrotic features, as indicated by *COL1A2*, *ALDOB*, and *HAVCR1* expression (Extended Data Fig. 3e)^7,44,45^. This pattern aligned with increased fibrosis and inflammation observed around injured tubules in our CODEX datasets.

To further investigate molecular signaling in these regions, we estimated spatially localized pathway activities using PROGENy (see Methods). Areas enriched in aFIB, FIB, and MES/VSMC exhibited elevated TGFβ signaling activity, which is a known driver of fibrosis (Fig. 3f). Similarly, immune cell-rich regions (LEUK) displayed increased NFκB and TNFα signaling activities, indicative of an inflammatory microenvironment (Extended Data Fig. 3f, *orange arrow*). The spatial distribution of EMT and ECM scores further highlighted a shift toward fibrotic cell types in DN kidneys (Extended Data Fig. 3g).

Taking our analysis further, we employed MISTy^46^ to explore how a cell’s behavior is influenced by its immediate neighbors and the broader tissue context. Evaluating influences both within a spot (intrinsic view, Extended Data Fig. 3h) and within 15 spots (extended neighborhood, Extended Data Fig. 3h), we discovered that endothelial and podocyte cells are highly interdependent within a spot, while aFIB strongly predict the presence of FIB and MES/VSMC—echoing the cellular compositions of Niches 8 and 11. Additionally, the abundance of PT cells correlated with the presence of iPT, LEUK, and aFIB in the extended neighborhood (mirroring Niches 10 and 13), underscoring how PT-rich areas are particularly vulnerable to injury, which in turn triggers immune responses and fibroblast activation^47,48^.

We next assessed the interdependencies between immune cells and major kidney cell types within a spot (Extended Data Fig. 3i, j). Our analysis revealed that the abundance of PT and iPT cells were best predicted by the presence of immune cells, suggesting a strong association between immune infiltration and PT injury. The colocalization of M2 macrophages and B cells with iPT cells aligned with their established roles in chronic injury and fibrosis (Extended Data Fig. 3k, l)^49–51^ . Finally, by analyzing the spatial interplay between the cell types and signaling pathways, we confirmed that TGFβ signaling predominantly associated with pro-fibrotic cell types, and NFκB and TNFα signaling predicted by immune cells (Extended Data Fig. 3m).

Altogether, our study reveals the molecular and spatial landscape of human DN and uncovers PT injury niches defined by injured tubular cells, M2 macrophage infiltration, and localized fibrotic signaling. These immune-tubular microenvironments are consistently observed across patients and may sustain tubular injury and drive fibrotic progression, yet the molecular regulators controlling tubular reprogramming within them remain undefined.

### 2.4 c-Jun upregulation results in PT injury

To identify candidate regulators of tubular reprogramming, we analyzed transcriptional programs along the PT injury continuum. Using publicly available sn/scRNA-seq datasets from DN samples, we reconstructed single-cell trajectories to map dynamic transitions from healthy to injured states.

This analysis revealed a continuum of PT cell states transitioning from homeostatic to injury-associated states (iPT) (Fig. 4a). This observation aligns with our cellular proportion analysis, which showed an increased abundance of iPT and a corresponding decrease of PT cells in DN kidneys compared to control kidneys (Extended Data Fig. 4a).

**Figure 4.**
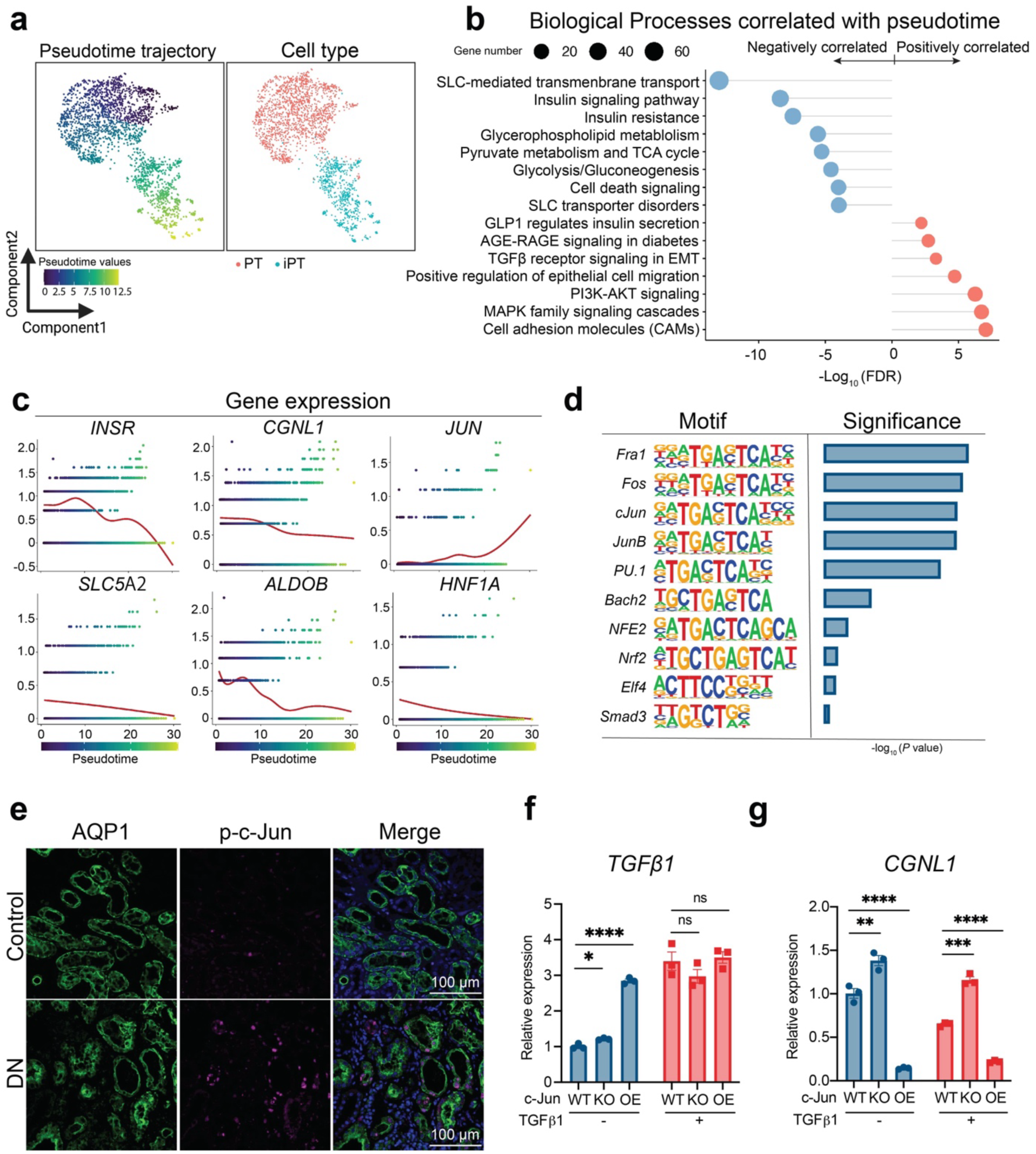
c-Jun is a central regulator in PT injury. a, Single cell trajectory analysis of PT and iPT cells. Cell types are shown in different colors. b, Pathway enrichment analysis for genes positively or negatively correlated with pseudotime trajectory. c, Prominent gene expression dynamics along pseudptime trajectory are shown. DN related genes (left, *INSR*, *SLC5*A2), epithelial marker genes (middle, *CGNL1*, *ALDOB*) and cell transition related transcription factors (right, *JUN*, *HNF1A*). d, Significance of enriched motifs and their corresponding transcription factors binding to these motifs along the pseudotime trajectory. e, Immunostaining of AQP1 and phosphorylated c-Jun (p-c-Jun) expression in DN and Control tissues. Scale bar, 100 µm. f and g, mRNA expression of *TGFβ1* and *CGNL1* in WT, c-Jun-KO and c-Jun-OE/HK2 cells with or without TGFβ1 treatment for 24 hours. Data are adjusted using unpaired two-tailed t-test. *P < 0.05, **P < 0.01, ***P < 0.001, ****P < 0.0001.

To gain mechanistic insights into this transition, we examined differentially expressed genes (DEGs) along the pseudotime trajectory. Genes downregulated during this transition were associated with diabetic kidney dysfunction, including pathways related to insulin resistance, glycerophospholipid metabolism, and SLC transporter disorders. In contrast, genes upregulated along the trajectory were enriched in pro-fibrotic pathways, including TGFβ receptor signaling, PI3K-AKT signaling and MAPK family signaling (Fig. 4b). Consistent trends were observed when comparing PTs with injured PTs (Extended Data Fig. 4b). Notably, renal function-related genes such as *INSR* and *SLC5A2* were significantly downregulated over pseudotime (Fig. 4c, *left*), consistent with previous findings of their reduced expression in diabetic kidney disease (DKD) samples^52,53^. Similarly, epithelial integrity markers *CGNL1* and *ALDOB* exhibited progressive downregulation (Fig. 4c, *middle*), supporting the notion of EMT-associated loss of epithelial characteristics^54,55^.

Further analysis of transcription factor dynamics identified two factors with opposing expression trends over pseudotime (Fig. 4c, *right*). *HNF1A*, a master regulator of PT differentiation and reabsorption function^56^, exhibited progressive downregulation, aligning with previous study that have shown PT dysfunction following HNF1A loss^57^. In contrast, *JUN* (which encodes the c-Jun protein and is part of the AP-1 complex involved in cell proliferation^58^) showed a marked increase in expression over pseudotime, indicating that c-Jun likely serves as a key regulator of PT injury.

Supporting this, we analyzed chromatin accessibility changes in PT injury using a published snATAC-seq dataset from DKD kidneys generated by the Humphreys lab^6^. This analysis revealed that AP-1 family motifs, including c-Jun binding motifs, were among the most enriched motifs in differentially accessible chromatin regions (DARs) associated with the PT injury (Fig. 4d, Extended Data Fig. 4c). We mapped these DARs to protein-coding genes and identified 723 genes overlapping with sn/scRNAseq DEGs, many of which were linked to NFκB signaling, insulin resistance, SLC-mediated transport, and TGFβ receptor signaling (Extended Data Fig. 4e), further indicating c-Jun’s role in PT injury.

To validate c-Jun’s involvement, we stained DN biopsies and observed active c-Jun (phospho-Ser73) in patient PTs (Fig. 4e). In cultured human kidney proximal tubular cells (HK2 cells), overexpression of c-Jun (c-Jun-OE) led to the upregulation of *TGFβ1* and *PAI-1*, both key mediators of fibrosis (Fig. 4f, Extended Data Fig. 4f, g), indicating that c-Jun activates TGFβ1 signaling. Furthermore, c-Jun-OE strongly repressed the expression of epithelial markers *CGNL1* and *CDH1* (Fig. 4g, Extended Data Fig. 4h), indicating that c-Jun promotes the loss of epithelial identity. Importantly, TGFβ1 treatment further repressed epithelial gene expression, whereas the knockout of c-Jun (c-Jun-KO) reversed the repression (Fig. 4g, Extended Data Fig. 4h).

To further assess *JUN* is functionally associated with the injured PT cell state, we performed an *in-silico JUN* knockout using CellOracle^59^. Our findings showed that knockout of *JUN* markedly disrupted the trajectory from PT to iPT states (Extended Data Fig. 4i, j) and significantly reduced the expression of multiple iPT related genes and related profibrotic pathways (Extended Data Fig. 4k-m), suggesting that knockout of c-Jun in PTs protect from fibrosis in DKD patients. These findings support a role for c-Jun as a transcriptional driver in PT injury.

### 2.5 STZ-induced diabetes is associated with c-Jun activation in proximal tubules

To address whether c-Jun is activated in a diabetic context *in vivo*, we employed a streptozotocin (STZ)-induced diabetes model in 11-week-old male C57BL/6 mice, administered at 50 mg/kg for five consecutive days (Fig. 5a).

**Figure 5.**
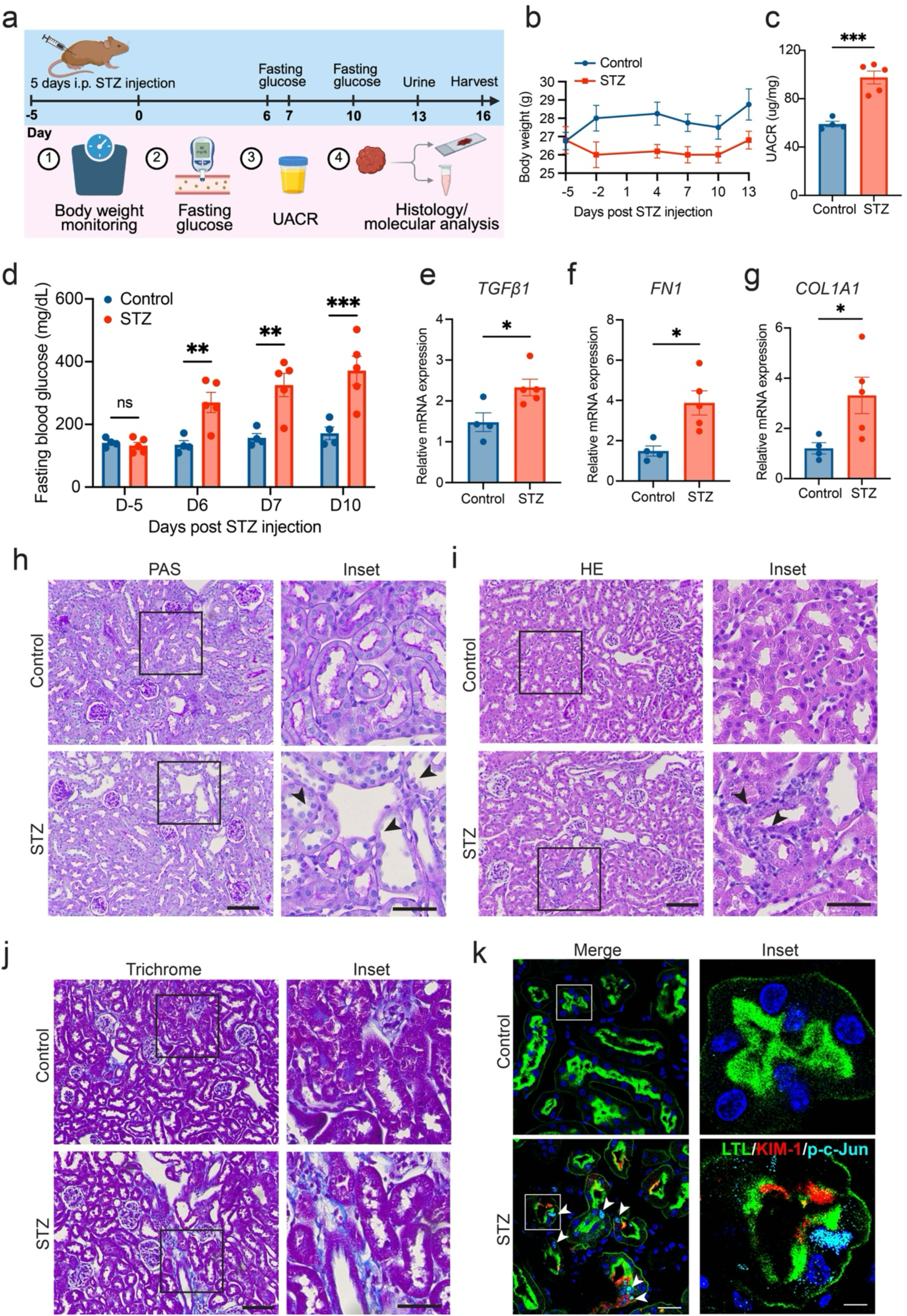
STZ-induced diabetic mice show hyperglycemia and increased c-Jun activation in proximal tubules. a. Schematic of the experimental design for STZ-induced diabetes in C57BL/6 male mice. b. Body weight curves showing weight changes in control and STZ-treated mice over time. c. Urine albumin-to-creatinine ratio (UACR) measured at day 13 post-STZ injection. d. Fasting blood glucose levels measured at the indicated time points following STZ administration. e–g. qPCR analysis of kidney tissue showing relative mRNA expression of *Tgfb1*, *Fn1*, and *Col1a1* in control (n = 4) and STZ-treated (n = 5) mice. Data are presented as mean ± SEM. h–j. Representative PAS, H&E and Trichrome staining showing tubular morphology in control and STZ-treated kidneys. Scale bar, 100 µm; inset, 50 µm. k. Representative immunostaining for LTL, KIM-1, and phosphorylated c-Jun (p–c-Jun) in control and STZ-treated kidney sections. Scale bar, 25 µm; inset, 5 µm.

STZ-treated mice developed sustained hyperglycemia, as shown by elevated fasting blood glucose levels across multiple time points (Fig. 5d). Body weight showed a slight decrease following STZ treatment but remained within ∼2 g of control mice throughout the experiment (Fig. 5b). Urinary albumin-to-creatinine ratio (UACR) was increased in STZ-treated mice at the time of collection (day 13 post-injection) (Fig. 5c). qPCR analysis revealed increased expression of *Tgfb1*, *Fn1*, and *Col1a1* in STZ-treated kidneys (Fig. 5e–g). These genes are associated with extracellular matrix production and fibrotic remodeling, indicating that transcriptional programs linked to fibrosis are already engaged at this stage.

Histological analysis showed tubular injury in STZ-treated kidneys. PAS staining revealed loss of brush border and tubular dilation in some tubules (*black arrows*, Fig. 5h), along with mild inflammatory cell infiltration (*black arrows*, Fig. 5i). Trichrome staining showed focal fibrotic deposition at this stage (Fig. 5j).

Immunofluorescence analysis demonstrated the activation of c-Jun (p–c-Jun) signal in injured proximal tubular cells in STZ-treated kidneys compared to controls (Fig. 5k), indicating that diabetic metabolic stress is sufficient to activate c-Jun signaling in proximal tubules in vivo. Together, these data show that c-Jun activation occurs in proximal tubules under diabetic conditions and coincides with tubular injury and early remodeling changes.

### 2.6 Tubular c-Jun activation induces chronic inflammation and kidney fibrosis

To assess the functional role of c-Jun *in vivo*, we established a tetracycline-inducible c-Jun mouse model (*c-Jun^tetO^ Pax8-rtTA)* by crossing *c-Jun^tetO^* mice^60^, which harbor a tetracycline-inducible c-Jun transgene, with Pax8-rtTA transgenic mice^61^, which express high levels of reverse tetracycline-dependent transactivator (rtTA) in renal tubules driven by the Pax8 promoter (Fig. 6a). Doxycycline (Dox) administration activated c-Jun expression specifically in renal tubules.

**Figure 6.**
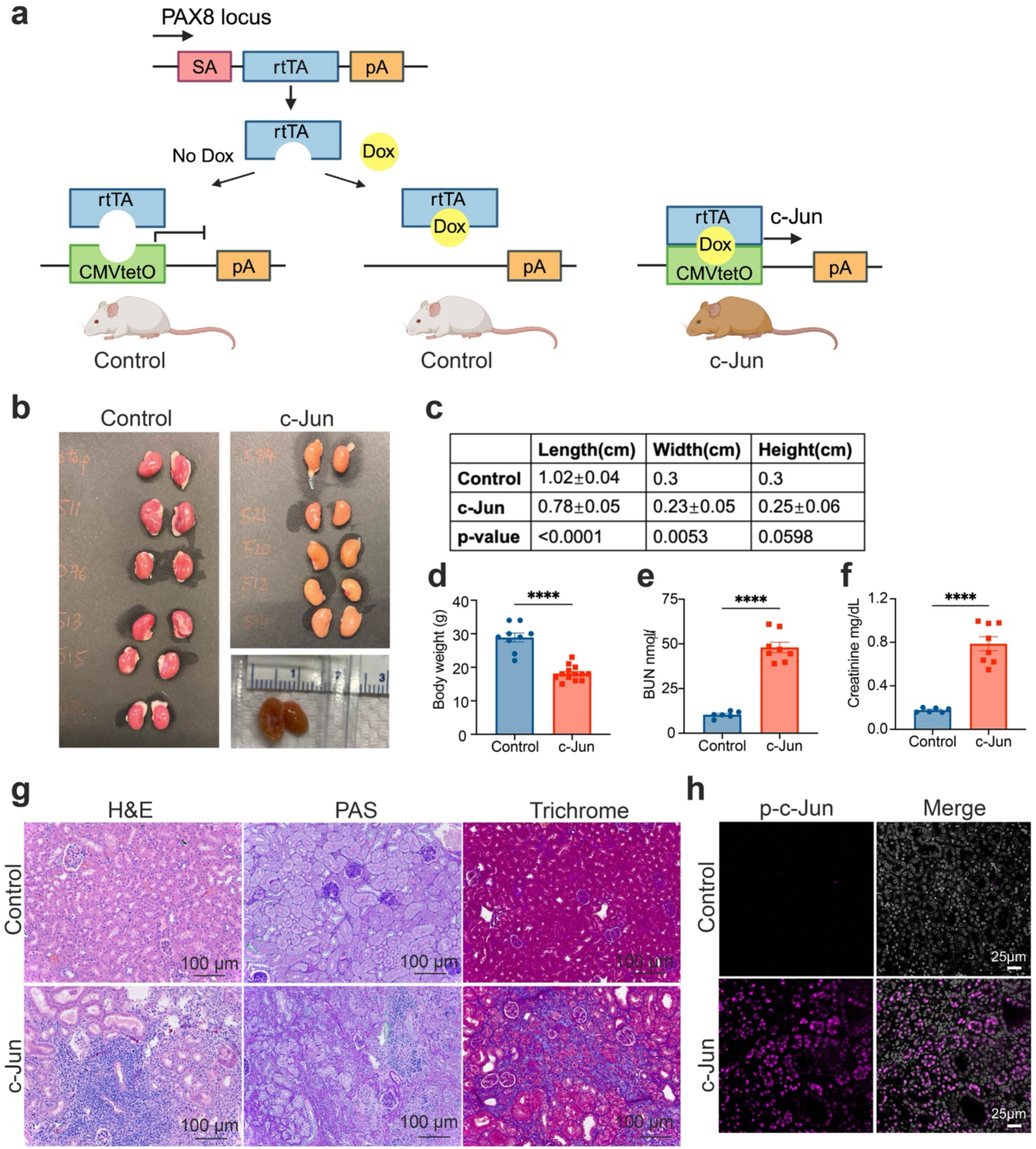
Characterization of tubule-specific c-Jun mouse model. a, Schematic of design of c-Jun mouse model. The construct at the PAX8 locus (rtTA) is coupled with the tetracycline operator minimal promoter (tetO) and leads to c-Jun overexpression specifically in renal tubules upon doxycycline (Dox) administration (right). The model without Dox (left) or have rtTA construct but lack c-JuntetO are served as control mice. b, Kidneys showing fibrosis developed after 12 weeks of Dox administration in *c-Jun^tetO^ Pax8-rtTA* mice. c, Kidney size changes in Control (n=9) and c-Jun mice (n=13), data are presented as mean with S.D. P-values are adjusted by t-test. d, Body weight changes in control (n=9) and c-Jun mice (n=13). e, Blood urea nitrogen (BUN) levels and f, Creatinine levels. Control, n=6. c-Jun, n=8. g, Representative images of H&E, PAS and Trichrome staining of kidneys from the indicated experimental groups. Scale bar, 100 µm. h, Representative images of immunostaining for phosphorylated c-Jun (p-c-Jun). Scale bar, 25 µm.

After 12 weeks of Dox treatment, both male and female c-Jun-overexpressing mice (c-Jun mice) exhibited marked renal impairment, including a significant reduction in kidney size and body weight loss compared to controls (Fig. 6b–d). Functional assays revealed elevated blood urea nitrogen (BUN) and creatinine levels, clear indicators of renal dysfunction (Fig. 6e, f).

Histological analysis confirmed tubular hypertrophy and atrophy, interstitial fibrosis, and immune cell infiltration in c-Jun mice (Fig. 6g, Extended Data Fig. 5a). Notably, these pathological changes were kidney-specific, as no abnormalities were observed in other organs, including the spleen, liver, lungs, or heart (Extended Data Fig. 5c).

Immunostaining confirmed c-Jun activation along with increased expression of the fibrotic markers (αSMA and FSP1) in c-Jun mice (Fig. 6h, Extended Data Fig. 5b). Additionally, cytokine profiling from serum and kidney tissue supernatants showed significantly increased levels of pro-inflammatory cytokines, including chemokines associated with macrophage recruitment (CCL3, CCL5, CXCL10)^62,63^ and M1–to–M2 macrophage transition (MIP2, CCL7)^64,65^(Extended Data Fig. 5d, e). These findings align with previous reports demonstrating c-Jun’s role in macrophage recruitment via direct transcriptional activation of *CCL5* and *CXCL10* in tumor models^66,67^.

To further explore the immune landscape in c-Jun-induced renal fibrosis, we employed mass cytometry (CyTOF)^68^ with a tailored antibody panel (Supplementary Table 3). Unsupervised clustering identified nine major cell types, including immune cells (B cells, CD4^+^ T cells, CD8^+^ T cells, Tregs, macrophages, and neutrophils), endothelial cells, epithelial cells, and CD45⁻CD31⁻CD324⁻ cells (likely fibroblasts and others) (Fig. 7a and b, Extended Data Fig. 6a). In c-Jun mice, we observed a loss of epithelial and structural cells, consistent with tubular damage (Fig. 7a, b, Extended Data Fig. 6b). Concurrently, CD4^+^ T cells, CD8^+^ T cells, Tregs, and macrophages were significantly increased, while neutrophils were significantly reduced (Fig. 7c, Extended Data Fig. 6b). Given that neutrophils are among the first responders during acute inflammation, their decline may suggest a transition toward a chronic inflammatory and tissue-repairing state^69,70^.

**Figure 7.**
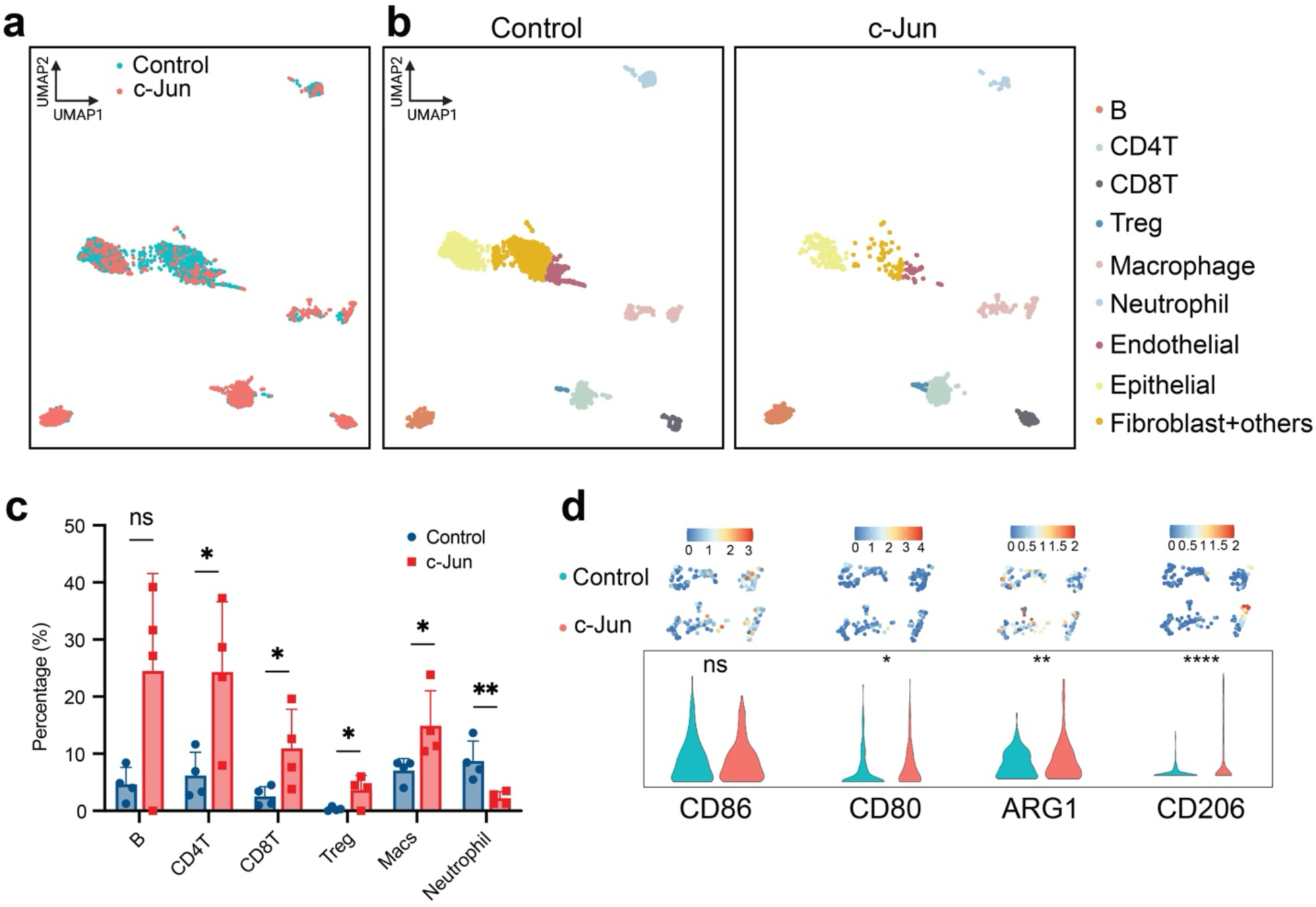
Immune phenotype characterization of c-Jun mice. a, UMAP plot of mass cytometry (CyTOF) dataset. b, UMAP plot of immune subpopulations in control and c-Jun mice. c, Percentage of CD45^+^ leukocytes (B cells, CD4^+^ T cells, CD8^+^ T cells, Tregs, macrophages, and neutrophils) in the indicated experimental groups are quantified by mass cytometry. d, Violin plots and corresponding deconvoluted abundance graphs depict M1 macrophages (CD86 and CD80 markers) and M2 macrophages (ARG1 and CD206 markers) in the kidneys of control and c-Jun mice. The ASINH ratio transformation was applied to normalize the marker expression intensities. Data from mass cytometry assays were obtained from 4 control and 4 c-Jun mice.

Consistently, we observed an increased abundance of M2 macrophages (marked by ARG1 and CD206), known for their role in tissue repair and ECM remodeling^71,72^ (Fig. 7d), while M1 macrophage activation (CD80 and CD86 markers), which dominate inflammation and tissue injury, was less pronounced. The macrophage profile in c-Jun mice mirrored the enrichment of CD68^+^ and CD163^+^CD68^+^ macrophages found in CN7 and CN8 of the iFME in DN patients (Fig. 2d). Additionally, we observed increased PD1 and PD-L1 expression in CD4^+^ and CD8^+^ T cells indicated T cell exhaustion likely due to sustained inflammation (Extended Data Fig. 6c).

### 2.7 Tubular c-Jun activation perturbs systemic metabolic homeostasis

To assess whether tubular c-Jun activation affects systemic glucose regulation, male *c-Jun^tetO^ Pax8-rtTA* mice were treated with Dox with or without high-fat diet (HFD) for 10 weeks (Extended Data Fig. 7a). c-Jun+HFD mice gained more weight from week 4 onward, but this increase was mitigated by c-Jun induction, resulting in comparable endpoint body weights to controls (Extended Data Fig. 7b, c).

Glucose tolerance tests (GTT) showed that c-Jun activation alone impaired glucose tolerance compared with control mice, indicating that tubular c-Jun activation is sufficient to disrupt systemic glucose homeostasis. The GTT response in c-Jun+HFD mice was similar to that of c-Jun mice, suggesting that HFD did not further worsen glucose intolerance (Extended Data Fig. 7d, e). Insulin tolerance tests (ITT) revealed an exaggerated hypoglycemic response in c-Jun mice compared with controls, whereas HFD feeding partially blunted this response, possibly reflecting an HFD-induced insulin resistance (Extended Data Fig. 7f, g).

The extent of injury and fibrosis in c-Jun+HFD mice was comparable to that in c-Jun mice, indicating that HFD did not further exacerbate the morphological changes (Extended Data Fig. 7h), although *JUN* expression was further increased in c-Jun+HFD mice compared with c-Jun only (Extended Data Fig. 7i), suggesting that metabolic stress amplifies c-Jun activation. qPCR analysis of kidney tissue revealed downregulation of gluconeogenic genes (*G6PC1* and *PCK1*) and the proximal tubular endocytic receptor *LRP2* in c-Jun mice compared with controls, indicating disrupted tubular metabolic and reabsorptive functions (Extended Data Fig. 7j-l).

These results suggest that c-Jun activation disrupts systemic glucose homeostasis and renal metabolic gene expression, while HFD exposure modifies the associated metabolic responses without substantially exacerbating renal fibrosis under the conditions tested.

### 2.8 SLC4A4 downregulation accompanies c-Jun-induced tubular injury

To characterize the metabolic alterations associated with c-Jun-induced tubular injury, we analyzed gene expression changes in injured proximal tubules. Pathway analysis revealed broad downregulation of solute carrier (SLC) family members, with SLC4A4, encoding the sodium bicarbonate cotransporter NBCe1, among the most prominently suppressed (Fig. 4b, Extended Data Fig. 4b,8a). Along the PT to iPT trajectory, SLC4A4 expression was significantly downregulated, accompanied by decreased chromatin accessibility at the locus (Extended Data Fig. 8b), suggesting potential disruption of bicarbonate homeostasis during tubular injury in DN.

Functional perturbation of SLC4A4 in human HK-2 cells confirmed its role in intracellular pH (pHi) regulation. SLC4A4-OE cells exhibited significantly higher pHi compared to SLC4A4-KO cells when exposed to HCO_3_^-^-buffered solution (pH 7.4), SLC4A4-KO cells displayed a decreased buffering capacity following cytosolic acidification by butyric acid (40 mM butyrate) (Extended Data Fig. 8c). These data suggested that SLC4A4 is vital for maintaining the optimal pHi of human proximal tubular cells by regulating intracellular bicarbonate levels.

In the context of tubular c-Jun activation, we observed that SLC4A4 expression was reduced in HK2 cells and markedly decreased in severely injured tubules of c-Jun-induced mice (Extended Data Fig. 8d-i). Together, these data indicate that SLC4A4 dysregulation accompanies c-Jun associated tubular injury and may contribute to impaired acid-base homeostasis in disease.

## 3 Discussion

Diabetic nephropathy (DN) is a progressive kidney disease characterized by both glomerular and tubulointerstitial injury, with fibrosis as a hallmark of advanced stages. While glomerular abnormalities have been extensively studied, accumulating evidence suggests that tubular injury plays a more active role in DN pathogenesis than previously recognized. However, the transcriptional and metabolic pathways linking tubular stress to fibrotic remodeling remain incompletely defined. This knowledge gap is largely due to the limited spatial and single-cell resolution of conventional transcriptomic and histological methods. We have addressed these gaps by employing an integrative, high-resolution multi-omics approach to dissect the cellular and molecular mechanisms governing DN progression.

By integrating multiplexed protein imaging, spatial transcriptomics, and single-cell/nucleus sequencing, we provide a multi-scale map of DN with emphasis on spatially organized cell states and microenvironments. To our knowledge, this is the first study to apply CODEX to human DN biopsies and define immune-fibrotic microenvironments in situ. Our analysis reveals disease-associated cellular niches enriched in injured tubules, activated fibroblasts, M2 macrophages, and extracellular matrix deposition. While inflammation and fibrosis are well-established features of DN, our data offer additional resolution by capturing their spatial co-localization and cellular composition at single-cell resolution. Building on this spatial framework, we prioritized transcriptional regulators associated with tubular injury and identified c-Jun as a compelling candidate.

A key contribution of our study is the demonstration that c-Jun directly drives tubular injury and fibrotic remodeling in the diabetic kidney. Integrated trajectory and chromatin accessibility analyses identified c-Jun as a top transcriptional regulator associated with PT injury. Functional perturbation in HK-2 cells showed that c-Jun activation induces pro-fibrotic gene expression while suppressing epithelial identity.

To determine whether c-Jun is activated under diabetic conditions in vivo, we employed an STZ-induced diabetic mouse model and observed increased phosphorylated c-Jun signal in proximal tubules, accompanied by tubular injury and early remodeling changes. These findings support the clinical relevance of c-Jun activation in the diabetic kidney. In parallel, we generated a tubule-specific inducible c-Jun mouse model to examine the functional consequences of sustained c-Jun activation. Activation of c-Jun in tubules induced tubular injury, nephron loss and progressive interstitial fibrosis, closely mirroring key pathological features of DN^73,74^. This was accompanied by persistent immune activation, characterized by upregulation of pro-inflammatory cytokines and infiltration of immune cells. These findings align with inflammatory signatures observed in DN patients^75,76^ and suggest that c-Jun activation promotes a feedforward immune-fibrotic loop, in which chronic inflammation and fibrosis reinforce each other and contribute to sustained tubular injury and disease progression.

In addition, tubule-specific c-Jun activation also perturbed systemic glucose homeostasis, as evidenced by impaired glucose tolerance, exaggerated insulin-induced hypoglycemia, and suppression of renal gluconeogenic genes. These findings indicate that c-Jun-induced tubular injury is accompanied by broader metabolic disturbances. Metabolic disturbances such as impaired glucose tolerance and susceptibility to hypoglycemia are well-recognized complications of CKD and DN, arising in part from reduced renal gluconeogenesis and altered insulin clearance^77–79^. In our c-Jun model, tubule-specific activation recapitulated several of these features, suggesting that tubular injury is sufficient to perturb systemic metabolism. We also observed reduced SLC4A4 expression in injured tubules, although the mechanistic relationship between c-Jun activation and SLC4A4 regulation remains to be determined.

Our study has several limitations. First, although our findings support the relevance and functional contribution of c-Jun activation in diabetic kidney injury, it remains unclear which c-Jun-associated transcriptional injury programs in DN are shared with, or distinct from those in other forms of CKD. Future comparative multi-omics profiling across diabetic and non-diabetic CKD samples, such as hypertensive nephropathy or FSGS, will be helpful in addressing this question. Second, while our inducible c-Jun model enables precise manipulation in kidney tubules, it does not distinguish among tubular subsegments. Future studies using segment-specific Cre drivers, such as *SLC34A1-Cre* developed by the Humphreys lab^80^, may help dissect the PT-specific effects of c-Jun activation and downstream metabolic alterations.

In conclusion, our study provides a spatially resolved, multi-omics framework for understanding DN pathogenesis and identifies c-Jun as a central transcriptional regulator of tubular injury and fibrosis. Using an inducible tubule-specific mouse model, we show that c-Jun activation drives chronic inflammation and progressive kidney fibrosis, while a diabetic model further supports the relevance of c-Jun activation in the injured diabetic kidney. Together, these findings establish c-Jun as a potential therapeutic target for diabetic kidney disease.

## 4 Methods

### 4.1 Samples and Ethical compliance

We have complied with all ethical regulations related to this study. All patients from Stanford Hospital who were included in the study gave consent to take part in the study with no participant compensation following Institutional Review Board (IRB) approval (IRB protocol 47382). Histologic sections were reviewed by a renal pathologist and laboratory data was abstracted from the medical record. All animal studies were performed in accordance with the Stanford University Institutional Animal Care and Use Committee and National Institutes of Health guidelines (APLAC#30911 and APLAC#30912). All mice were housed in the Stanford University SIM1 Animal Facility (Stanford Institute for Medicine 1) and maintained in specific pathogen-free conditions.

### 4.2 CODEX assay

#### 4.2.1 Construction of tissue microarray (TMA)

FFPE tissue blocks were retrieved from the tissue archive at the Stanford Health Care Department of Pathology. 66 unique different tissues were selected (3 tissue regions of 20 diabetic nephropathy patients and 2 healthy donors; for details see Supplementary Table 1. The fibrosis and normal tissue regions were annotated on corresponding hematoxylin and eosin (H&E)-stained sections by a board-certified pathologist (N.A.B). The TMA was constructed from 1-mm-diameter cores punched from the tissue blocks.

#### 4.2.2 CODEX image acquisition and segmentation

Multiplexed CODEX analysis was performed using а panel of antibodies (Extended Data Fig. 1a, b) conjugated to custom DNA barcodes and detector oligos and common buffers, with a robotic imaging setup, according to the instructions for CODEX staining of FFPE specimens from Akoya Biosciences (https://www.akoyabio.com/). All steps followed the Akoya CODEX instructions. Acquired images were preprocessed (alignment and deconvolution) and segmented with HALO software (https://indicalab.com/).

#### 4.2.3 CODEX quantification and Cellular neighborhood (CN) analysis

We implemented a deconvolution approach “Celesta”^30^ to assign the most likely cell type to each cell in the presence of cell-type mixtures (Supplementary Table 5). As such, for each mixed cell type group, we extracted a feature intensity submatrix consisting of those cells and the features relevant to the cell type mixtures they represented. We used the identify neighborhood’s function (method = “hierarchical”, min_neighhorhood_size = 10, radius = 50 for DN samples, min_neighhorhood_size = 20, radius = 50 for Control samples) in the SPIAT (V0.4) package^34^ of R software to identify CNs of each tissue. Cells of interest were those we identified and annotated in the study. While cells that could not be annotated were not analyzed. A R script was designed to analyze the proportion and number of various cell types in each CN. We then applied the ggplot function in the ggplot2 package (V 3.3.5) of R to plot the spatial distribution of cells and CNs. The mixing_score_summary function in the SPIAT was used to calculate the mix score and the calculate_minimum_distances_between_celltypes function in the SPIAT was used to calculate the minimum cell distances of each sample.

In addition to the methods listed below, the specific details of statistical tests (values of n, p values, type of statistical test, definition of center etc.) are reported in the figure legends.

### 4.3 Spatial gene expression assay (Visium)

#### 4.3.1 Spatial transcriptomics method

FFPE tissue blocks of 4 diabetic nephropathy patients and 4 healthy donors were selected for Visium assay; for details see Supplementary Table 2. The RNA quality of the human kidney FFPE sample was checked by the Quick-RNA FFPE Kit (Zymo Research, R1008) according to manufacturer’s protocol. RNA quality was examined using an Agilent bioanalyzer, and samples with DV200 > 50% were selected. Then a 5 µm tissue sample was cut onto the 10× Visium Spatial Gene Expression Slide according to manufacturer’s protocol (10X Genomics, CG000408). After deparaffinization, H&E staining was performed (10X Genomics, CG000409). We used Keyence BZ-9000 microscope for whole-slide imaging. After scanning, de-crosslinking, probe hybridization, probe release and extension, library preparation was performed using Dual Index Kit TS Set A (10X Genomics, PN-1000251). Quality control for the libraries was performed using the Agilent Bioanalyzer High Sensitivity DNA Assay. Libraries were sequenced using the Illumina Hiseq 4000 system with 2 × 150 paired-end kits. Demultiplexing used 28 bp Read1 (cell barcode and UMI), 10 bp i7, 10 bp i5, and 50 bp Read2 (transcript).

#### 4.3.2 Spatial transcriptomics data analysis

Filtered feature-barcode expression matrices from SpaceRanger (v1.3.2) were used as initial input for the spatial transcriptomics analysis using Seurat (v4.0.1). Spots with less than 200 measured genes and less than 3 UMIs were filtered out. Individual count matrices were normalized with SCTransform^81^, and additional log-normalized (size factor = 10,000) and scaled matrices were calculated for comparative analyses using default settings. Cell-type compositions were calculated for each spot using Seurat. Reference expression signatures of major cell types were estimated using regularized negative binomial regressions and our integrated sn/scRNA-seq atlas. For each spot, we estimated signaling pathway activities with PROGENy’s^82,83^ (v1.12.0) model matrix using the top 1,000 genes of each transcriptional footprint and the SCTransform normalized data. We used MISTy’s^46^ implementation in mistyR (v1.2.1) to estimate the importance of the abundance of each major cell type in explaining the abundance of the other major cell types. Cell-type estimations of all slides were modelled in a multi-view model using three different spatial contexts: (1) an intrinsic view that measures the relationships between the deconvolution estimations within a spot, (2) a juxta view that sums the observed deconvolution estimations of immediate neighbors (largest distance threshold = 5), and (3) a para view that weights the deconvolution estimations of more distant neighbors of each cell type (effective radius = 15 spots).

To identify groups of spots in the different samples that shared similar cell-type compositions, we transformed the estimated cell-type proportions of each spatial transcriptomics spot and slid into isometric log ratios, and clustered spots into groups. These niches represent groups of spots that are similar in cell composition and represent potential shared structural building blocks of our different slides. To complement the repertoire of niches identified with cell-type compositions, we integrated and clustered the Visium spots of all slides using their log-normalized gene expression. We called these clusters molecular niches. Integration and clustering of spots was performed with the same methodology as the one used to create the snRNA-Seq atlas.

#### 4.3.3 Quantification of ECM and EMT scores

To quantify the ECM and EMT scores, we used the ‘AddModuleScore’ function from the Seurat R package. This function calculates an average expression score for a user-defined gene set in each cell, adjusted by control gene sets matched for average expression levels, thereby generating a relative activity score per cell. For ECM score, we utilized gene sets curated in a previously published ECM gene resource^42^. Specifically, we included 1,027 genes encoding core ECM components, along with a subset of 274 genes representing collagens, glycoproteins, and proteoglycans. For EMT score, we used the EMT gene list defined in the EMTome database^43^, which provides a comprehensive compilation of EMT-related genes. All genes listed in EMTome were included in our analysis.

### 4.4 Single cell RNA-seq data analysis

#### 4.4.1 Differential expression analysis

Here we collected two published single cell RNAseq datasets (GSE195460 and GSE211785). The Seurat V4 data integration pipeline was used to batch correct the data through the canonical correlation analysis (CCA) method. According to a benchmark comparison study conducted by Tran and colleagues^84^, Seurat CCA was identified as one of the top three preferred batch integration techniques for this type of data. The R package SCTransform^81^ was used to normalize gene expression for each cell by fitting the Gamma-Poisson generalized linear model. The resulting log-transformed, normalized single-cell expression values were used for visualizations and differential expression tests. Statistically significant principal components were determined by a resampling test and were retained for the Uniform Manifold Approximation and Projection (UMAP) analysis. Differential expression analysis among clusters was conducted using a likelihood-ratio test, comparing cells within each cluster against all other cells. Gene A was defined as a biomarker for cluster X if it was detected in at least 25% of cells, had an adjusted p-value less than 0.05, and had a log e fold change of at least 0.25 between cells of cluster X and all other cells. These analyses were performed using the Seurat package v4.0. DEGs were analyzed for GO terms and KEGG pathways enrichment by using KOBAS^85^. A significance threshold of FDR < 0.05 was used during the enrichment analysis to identify significant results.

#### 4.4.2 Pseudotime trajectory analysis

Cell developmental trajectories were inferred using Monocle2 (version 2.99.3)^86^ with default parameters as recommended by the developers. Firstly, integrated gene expression matrices from specific cell type were exported from Seurat into Monocle to construct a CellDataSet. Secondly, the ‘setOrderingFilter’ function was applied to sort cells with the variable genes identified by the function of ‘differentialGeneTest’ (cutoff of q < 0.001). Finally, after dimensionality reduction using the ‘reduceDimension’ function (using the ‘DDRTree’ reduction method), a series of representative key role genes were revealed along the differentiation progress by the ‘plot_pseudotime_heatmap’ function.

#### 4.4.3 In-silico perturbation

Single-cell perturbation analysis of *JUN* was performed using celloracle^59^ on proximal tubule (PT) and injured proximal tubule (iPT) cells derived from our processed kidney single-cell RNA-seq dataset. *In-silico JUN* knockout was simulated using celloracle’s default perturbation module by computationally setting *JUN* activity to zero and propagating its effects through the inferred gene regulatory network. Predicted post-perturbation expression states were projected onto the original UMAP space, and changes in *JUN* target genes, and pathway signatures were quantified by comparing simulated *JUN* knockout profiles with the unperturbed baseline.

### 4.5 Single cell ATAC-seq data analysis

Published dataset of GSE195460 was collected. To control the data quality, low-quality cells were filtered out based on transcription start site (TSS) enrichment and the number of unique fragments. All peaks were merged to create a union peak set of which each peak was annotated as intergenic, promoter, exonic and intronic. For dimensionality reduction, UMAP was used to generate a 2D embedding for visualization by Signac^87^ cells were clustered using the Leiden algorithm. To annotate the clusters, a gene activity score matrix was created using the function addGeneScoreMatrix and marker genes were detected for each cluster using the function getMarkerFeatures. The same markers from snRNA-seq data were used to annotate the clusters. Correlation coefficient values were calculated between peaks and pseudotime. HOMER^88^ findMotifsGenome.pl was used to investigate the motif enrichment of pseudotime positive related peaks using default parameters.

### 4.6 Animal model of renal fibrosis

To generate doxycycline-inducible *c-Jun* transgenic mouse (*c-Jun^tetO^ PAX8-rtTA*), we crossed tetracycline-responsive Jun mice (*c-Jun^tetO^*)^60^ with transgenic Pax8-rtTA mice obtained from Jackson Laboratory (RRID: IMSR_JAX:007176). *c-Jun^tetO^ PAX8-rtTA* double-transgenic mice were healthy, with no detectable phenotype. 8-week-old male and female mice were used for all experiments. The sample size was chosen on the basis of our previous experience and for each experiment it is indicated in the figure legend. When treated with doxycycline, however, the homozygous mice only survived 10 days after administration. Therefore, heterozygous mice were selected to mimic of adult onset of chronic kidney disease by administrated with Dox at a concentration of 1mg/ml in drinking water for 12 weeks. Investigators were not blinded for group allocation but were blinded for the assessment of the phenotypic outcome assessed by histological analyses.

For STZ-induced diabetic mouse model, 11 weeks old male C57BL/6J mice were used. Diabetes was induced by intraperitoneal injection of streptozotocin (STZ; Cayman Chemical, catalog number 13104) at 50 mg/kg body weight once daily for five consecutive days. Mice were fasted for 4 hours prior to each STZ injection. STZ was freshly dissolved in citrate buffer (0.1 M, pH 4.5) prior to injection. Control mice received equivalent volumes of citrate buffer. Following STZ administration, mice were provided with 10% sucrose water to prevent acute hypoglycemia.

For metabolic study, 5-week-old male *c-Jun^tetO^ Pax8-rtTA* mice were treated with Dox with or without high-fat diet for 10 weeks.

#### 4.6.1 Renal function analysis

Blood was collected by retro-orbital sampling for measurement of BUN and creatinine. Serum BUN levels were measured using a Urea Assay Kit (Sigma-Aldrich, MAK006). Serum creatinine levels were measured using a Mouse Creatinine Assay Kit (Crystal Chem, 80350), according to the manufacturers’ instructions.

Fasting blood glucose levels were measured at the indicated time points following STZ administration using a glucometer (True METRIX, McKesson, USA). Mice were fasted for 4 hours prior to each measurement.

For urine albumin-to-creatinine ratio (UACR), urine samples were collected at day 13 post-STZ injection. Urinary albumin levels were measured using a Mouse Albumin ELISA Kit (Bethyl Laboratories, E99-134), and urinary creatinine levels were measured using a Mouse Creatinine

Assay Kit (Crystal Chem, 80350), according to the manufacturers’ instructions. UACR was calculated for each sample.

For GTTs, mice were fasted for 4 hours and injected intraperitoneally with glucose (2 g/kg body weight). Blood samples were collected from the tail vein at 0, 15, 30, 60, 90, and 120 minutes, and glucose levels were measured using a glucometer (True METRIX, McKesson, USA).

For ITTs, mice were fasted for 4 hours and injected intraperitoneally with insulin (0.75 U/kg body weight). Blood glucose levels were measured at 0, 15, 30, 60, 90, and 120 minutes.

#### 4.6.2 Histological analysis

Tissues of a proper size were fixed in 4% paraformaldehyde for 24 hours and dehydrated in 70% ethanol. Samples were then immersed in xylene and embedded in paraffin. Paraffin blocks were cut into 4μm. Slides were cleared in xylene and rehydrated before staining with either hematoxylin and eosin, Periodic acid-Schiff (PAS) or with Masson’s Trichrome according to the manufacturer’s recommended procedure. All images were captured on Nikon ECLIPSE Ni-E microscope. Collagen fibers were stained blue with a cord-like structure in Masson trichrome staining and collagen volume fraction was calculated using the HALO software.

#### 4.6.3 Immunofluorescence staining

For formalin-fixed paraffin-embedded (FFPE) sections from human DN samples and mouse kidneys, sections were deparaffinized, rehydrated, and subjected to antigen retrieval using Citrate buffer, pH 6.0 (Sigma-Aldrich, C9999-1000ml). Sections were blocked with 3% donkey serum in TBST buffer (TBS containing 0.05% Tween-20) and incubated with primary antibodies overnight at 4 °C. The following primary antibodies were used: goat anti–α-smooth muscle actin (α-SMA; 1:200; Novus Biologicals, NB300-978); rabbit anti–FSP1/S100A4 (1:200; Millipore Sigma, 07-2274); rabbit anti–phospho–c-Jun (Ser73) (1:200; Cell Signaling Technology, 3270S); rabbit anti–SLC4A4 (1:100; Novus Biologicals, NBP2-94138); mouse anti–AQP1 (1:100; Novus Biologicals, NB600-749). After washing with TBST buffer, sections were incubated with species-specific secondary antibodies conjugated to Alexa Fluor 488 or 647 (1:1000; Thermo Fisher Scientific) for 1 hour at room temperature. Nuclei were counterstained with DAPI for 5 minutes. Slides were mounted and imaged using a Leica SP8 confocal microscope (Leica Microsystems).

For mouse kidneys from the STZ-induced diabetes model, tissues were fixed in 4% paraformaldehyde for 1 hour, cryoprotected in 20% sucrose overnight at 4 °C, and embedded in OCT compound. Frozen sections were prepared for immunofluorescence staining. Sections were incubated with the following primary antibodies: goat anti–KIM-1 (1:200; R&D Systems, AF1817-SP); rabbit anti–phospho–c-Jun (Ser73) (1:200; Cell Signaling Technology, 3270S); Lotus tetragonolobus lectin (LTL; FITC-conjugated, 1:400; Invitrogen) was used to label proximal tubules. After washing, sections were incubated with appropriate species-specific secondary antibodies (1:400; Thermo Fisher Scientific) for 1 hour at room temperature. Nuclei were counterstained with DAPI, and images were acquired using a Leica SP8 confocal microscope (Leica Microsystems).

#### 4.6.4 Cytokine array (Luminex)

75μl of serum and kidney homogenate culture supernatant were collected for cytokines and chemokines measurement. This assay was performed by the Human Immune Monitoring Center at Stanford University. Mouse 48 plex Procarta kits (EPX480-20834-901) were purchased from Thermo-Fisher/Life Technologies, Santa Clara, California, USA, and used according to the manufacturer’s recommendations with modifications as described. Briefly: Beads were added to a 96 well plate and washed in a BioTek ELx405 washer. Samples were added to the plate containing the mixed antibody-linked beads and incubated overnight at 4°C with shaking. Cold (4oC) and Room temperature incubation steps were performed on an orbital shaker at 500-600 rpm. Following the overnight incubation plates were washed in a BioTek ELx405 washer and biotinylated detection antibody added for 60 minutes at room temperature with shaking. Plate was washed as described and streptavidin-PE was added for 30 minutes at room temperature.

Plate was washed as above and reading buffer was added to the wells. Each sample was measured in duplicate wells. Plates were read using a Luminex 200 or a FM3D FlexMap instrument with a lower bound of 50 beads per sample per cytokine. Custom Assay Chex control beads were purchased from Radix BioSolutions, Georgetown, Texas; and added to all wells.

#### 4.6.5 Single-cell mass cytometry (CyTOF)

Samples were processed following previously established protocols^89^. Briefly, cell samples were first fixed with 2% paraformaldehyde (PFA) at room temperature for 20 minutes and then washed twice with PBS containing 0.5% bovine serum albumin (BSA). To identify dead cells, the samples were stained with cisplatin (Sigma) at a final concentration of 25 μM for 1 minute at room temperature. The cisplatin staining was quenched by adding RPMI medium with 10% fetal bovine serum (FBS) for 3 minutes, and the cells were washed three times with CSM (PBS with 0.5% protease-free BSA and 0.02% NaN₃).

Afterward, samples were stained with metal-conjugated antibodies against cell surface markers. To prevent antibody aggregates, the surface antibody cocktail in CSM was filtered through a pre-wetted 0.1-µm spin column (Millipore) and incubated with the cells at room temperature for 1 hour. The cells were then washed with CSM.

For intracellular staining, cells were permeabilized using ice-cold methanol for 15 minutes and washed twice with CSM to remove residual methanol. Metal-conjugated antibodies targeting intracellular molecules were added, and the cells were incubated for 1 hour at room temperature. After washing again with CSM, the cells were incubated with an iridium-containing DNA intercalator (Cell-ID Intercalator-Ir, Fluidigm) at a final concentration of 125 nM in 4% PFA diluted in PBS for 20 minutes at room temperature to stain DNA.

Following intercalation and fixation, cells were washed with CSM. The cells were filtered and suspended in a solution containing EQ Four Element Calibration Beads (Fluidigm) before being analyzed using a CyTOF mass cytometer (Fluidigm). Post-measurement, individual files were processed by gating out doublets, debris, and dead cells based on cell length, DNA content, and cisplatin staining.

All data visualizations and statistical analyses were performed using a combination of GraphPad Prism (10.2) and R (4.4.1) software. Bar plots, including means and error bars (e.g., standard error or standard deviation), were generated using GraphPad Prism to display group-wise comparisons of key metrics. For more complex data visualizations, such as violin plots, pie charts, and Uniform Manifold Approximation and Projection (UMAP) plots, the R package ggplot2 was utilized. Violin plots were specifically generated to show the distribution and density of CyTOF ASINH-transformed values, providing insight into the distribution of marker expression across different groups. Pie charts were employed for categorical data representation, while UMAP plots were generated to visualize high-dimensional single-cell data, enabling the identification of clusters and patterns within the datasets.

#### 4.6.6 Mouse kidney tissue microarray

Paraffin-embedded, formalin-fixed kidney tissue from c-Jun and control mice were selected. Areas were randomly selected per sample and two 2-mm cores were taken from each kidney sample. Samples were processed following the same protocol as the TMA for CODEX assay.

#### 4.6.7 Quantitative RT-PCR analysis of mouse kidney tissue

Mouse kidney tissues (around 15mg) are homogenized and followed by RNA extraction according to the manufacturer’s instructions using the RNeasy Mini Kit (Qiagen). 0.5 μg of total RNA was reverse transcribed with Iscript™ Advanced cDNA Synthesis Kit (Bio-Rad, 1725038) and qRT-PCR was carried out on a QuantStudio 6 and 7 Pro Real-Time PCR System using the SYBR Green method (Applied Biosystems). RNA expression was calculated using the comparative Ct method normalized to GAPDH. Data were presented as 2^−(ΔΔCt)^ relative expression. The mouse primers used are listed in Supplementary Table 4.

### 4.7 Cell lines

#### 4.7.1 Construction of c-Jun/SLC4A4 overexpression and knockout HK2 cell lines

HK-2 cells were purchased from the American Type Cell Collection (ATCC, CRL2190). HK-2 cells were maintained in DMEM/F12, GlutaMAX Supplement (ThermoFisher, 10565018) supplemented with 2.5% FBS, 1% penicillin/streptomycin (Sigma-Aldrich, P0781) and 1% insulin-transferrine-selenium (ITS) (Sigma-Aldrich, I3146-5ML). The human c-Jun tet-on overexpression plasmid and CRISPR/Cas9-mediated c-Jun knockout plasmid were constructed previously^82^. The human lentiviral overexpression plasmid of SLC4A4 (pLV-C-GFPSpark-SLC4A4) was obtained from Sino Biological (HG18001-ACGLN). The human sgRNA: CRISPR-Cas9 vector of SLC4A4 was obtained from VectorBuilder (VB900145-2250khq).

Lentiviral particles were prepared by transfecting HEK293T cells with lentivectors, pMD2.G and psPAX2, with Lipofectamine 3000 (Thermo Fisher Scientific). Cell infection was performed as described previously^90^, followed by GFP or puromycin selection.

#### 4.7.2 TGF-β1 treatment

For c-Jun overexpression (HK2-c-JunOE) cell line, Dox (1 μg/ml, Sigma-Aldrich, D9891-10G) was applied 24 hours before TGF-β1 treatment to turn on c-Jun overexpression. Recombinant human TGF-β1 (10 ng/ml, PeproTech, 100-21C) was added to HK2-c-JunOE and c-Jun knockout (HK2-c-JunKO) cells for different time points. For inhibitor treatment, TGF-β Receptor Inhibitor LY2109761 (2.5μM, MedChem Express, HY-12075) was added to culture wells the same time as adding TGF-β. All experiments were performed in triplicates.

#### 4.7.3 Quantitative RT-PCR

Cell pellets were harvested and washed with PBS followed by RNA extraction according to the manufacturer’s instructions using the RNeasy Mini Kit (Qiagen). 1 μg of total RNA was reverse transcribed with Iscript™ Advanced cDNA Synthesis Kit (Bio-Rad, 1725038) and qRT-PCR was carried out on a QuantStudio 6 and 7 Pro Real-Time PCR System using the SYBR Green method (Applied Biosystems). RNA expression was calculated using the comparative Ct method normalized to GAPDH/ACTB. Data were presented as 2^−(ΔΔCt)^ relative expression. The primers used are listed in Supplementary Table 4.

#### 4.7.4 Western blot

For total protein preparation, cell lysates were prepared by RIPA Lysis Buffer (EMD Millipore, 20-188) with Complete™ Protease Inhibitor Cocktail (Roche, 11697498001). For membrane protein preparation, membrane proteins were extracted using Pierce™ Mem-PER™ Plus Membrane Protein Extraction Kit (Thermo Scientific, 89842). The protein concentrations were quantified using Bradford protein assay (Bio-Rad, 5000205). The protein lysates were heated for 15 min at 65°C in Laemmli SDS Sample Loading Buffer (Bioworld, 10570021-1) and loaded into 8% Bis-Tris Plus Gels (Thermo Scientific, NW00080BOX). Afterwards samples were transferred onto Nitrocellulise membranes (Thermo Scientific, LC2000) and the blots were probed with following primary antibodies overnight at 4°C : rabbit anti- Phospho c-Jun (Ser73) (D47G9) mAb (pJun; 1:1000; Cell Signaling Technology, 3270S), rabbit anti-c-Jun mAb (60A8) (c-Jun, 1:1000; Cell Signaling Technology, 9165T), rabbit anti-SLC4A4 antibody (SLC4A4, 1:1000, Novus Biologicals, NBP2-94138), rabbit anti-Vinculin (E1E9V) mAb (Vinculin, 1:1000, Cell Signaling Technology, 13901S), mouse anti- GAPDH (GT239) mAb (GAPDH, 1:1000, GeneTex, GTX627408), followed by incubation with HRP conjugated anti-mouse or anti-rabbit secondary antibody (Thermo Scientific) for 1 hour after washing and developed using WesternBright ECL-HRP Substrate (Advansta, K-12045-D20). Images were acquired by Chemidoc MP (Bio-Rad) and analyzed by ImageLab (Bio-Rad).

#### 4.7.5 Immunofluorescence staining

For staining surface SLC4A4 on HK2-c-Jun cell lines, 2 ξ 10^5^ cells were placed directly to glass bottom 30mm ξ 10mm culture dishes and incubated overnight. Cells were washed with PBS and blocked with a blocking buffer (1% BSA in PBS) for 1 hour. Cells were incubated with primary antibody rabbit anti-SLC4A4 antibody (SLC4A4, 1:100, Novus Biologicals, NBP2-94138) on ice for 2 hours. After washing with PBS, cells were incubated with 488 fluorophore-conjugated goat anti-rabbit secondary antibody (1:1000; Thermo Scientific, A11034) on ice for 1 hour. Cells were observed immediately using a Leica SP8 inverted confocal microscope.

#### 4.7.6 Intracellular pH measurement

To measure the intracellular pH (pHi) in HK2-SLC4A4 cell lines, a Leica SP8 inverted confocal imaging system and an acetoxymethyl ester pH-sensitive, fluorescent dye SNARF^TM^-5F (ThemoFisher Scientific, S23923) were used. The dye was loaded into the cells by incubating them with 3 μM SNARF^TM^-5F in HEPES-buffered solution (140 mM NaCl, 5 mM KCl, 10 mM D-glucose, 0.5 mM NaH_2_PO_4_, 10 mM HEPES, 1 mM MgCl_2_, and 2 mM CaCl_2_ set to pH 7.4) for 15 min at room temperature. Cells were then mounted on an open chamber of the Leica SP8 confocal microscope and the base pHi was recorded under HEPES-buffered solution. The standard HCO3^-^-buffered solution consisted of 114 mM NaCl, 5 mM KCl, 0.5 mM NaH_2_PO_4_, 10 mM D-glucose, 26 mM NaHCO_3_, 1 mM MgCl_2_, and 2 mM CaCl_2_ set to pH 7.4). For intracellular acidification, cells were challenged with 40 mM butyrate in HCO3^-^-buffered solution adjusted to pH 7.4. SNARF^TM^-5F was excited at 543 nm and the changes in fluorescence emission were collected in the bandwidths 580-620 nm and 630-670 nm. Images were obtained every 10 s with a 40× water-immersion objective. pHi was determined by using the ratio of fluorescence intensities from the dye at two emission wavelengths. The resulting ratios were converted into pHi values using a nigericin-based calibration technique. Specifically, cells were treated with commercial calibration solutions (Spexyte™ Intracellular pH Calibration Buffer Kit, Abcam, ab284662) corresponding to pH 6.0, 6.5, 7.0, 7.5 and 8.0. All experiments were performed at room temperature.

### 4.8 Visualization, statistics, and reproducibility

Data are presented as mean ± S.E.M if not specified otherwise in the legends. Unless otherwise stated, independent sample t test was used to compare the continuous variable in two groups and one-way ANOVA for multigroup comparisons (n.s. non-significant; P > 0.05; *P < 0.05; **P < 0.01; ***P < 0.001; ****P < 0.0001). Statistical analyses were performed using GraphPad Prism 10 (GraphPad Software) or as described in the Methods above. Results are presented in dot plots, with dots representing individual values. The number of samples for each group was chosen based on the expected levels of variation and consistency. The presented immunofluorescence micrographs and western blot images are representative. All schematic diagrams were created using BioRender.com under an academic license.

## 5 Data availability

All data associated with this study are present in the paper or the Supplementary Materials. The spatial data have been deposited in GEO under accession number GSE249740. Source data are provided with this paper.

## Acknowledgements

This work was supported by Dr. G. Wernig’s funds Startup Funds Pathology, Scleroderma Research Foundation, Ludwig Foundation and Stanford Diabetes Research Center (SDRC) Pilot and Feasibility Program (NIH P30DK116074). We thank Deana Colburg (Stanford University) for help with tissue microarrays, Dr. Weiruo Zhang (Stanford University) for help with the CELESTA algorithm, Dr. Jun Wu (Stanford University) and Pauline Chu (Stanford University) for frozen tissue sectioning and Dr. Lee Fradkin (Stanford University) for critical review of the manuscript. Graphical illustrations were created with BioRender.com (license to Q.D.) under an academic license. ChatGPT was used to refine the language of the manuscript.

## 6 Author contributions

G.W. and Q.D. conceived and designed the study. Q.D. and Y.L. conducted experiments, collected and analyzed the data and generated the figures, with contributions of Y.C.W., P.A.W., M.A., M.S., N.A.B. and V.C. provided clinical, pathological and immunological renal expertise, tissue sample collection for study cohort and performed CODEX imaging experiment. Q.D., Y.L. and G.W. wrote the manuscript with contributions from N.A.B., Y.C.W, S.Y.C, J.C.W. and V.C. All authors contributed to and approved the final version of the manuscript.

---

**Extended Data Figure 1.**
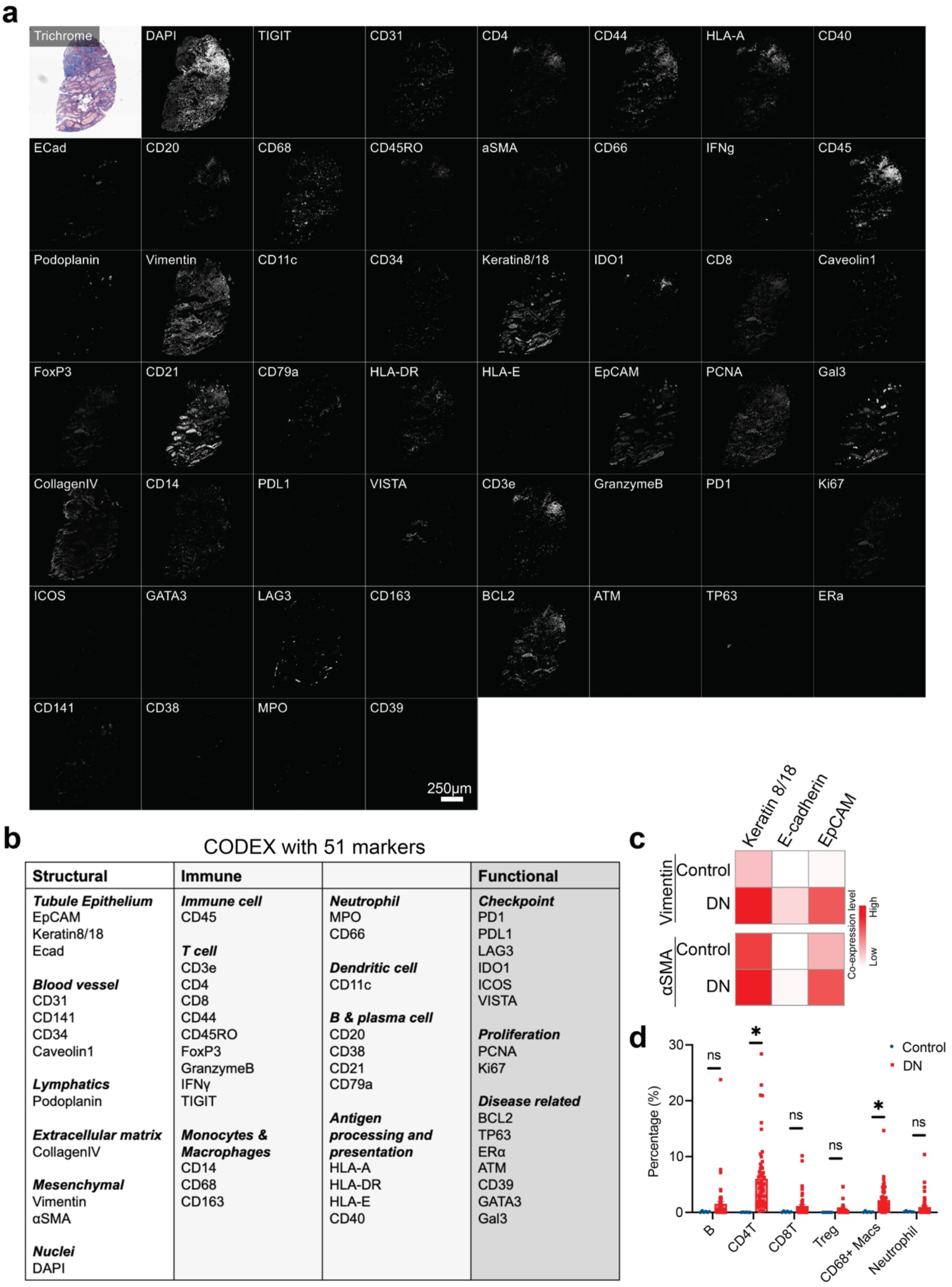
CODEX antibody panel used in the study. a, Each marker of the CODEX panel is shown individually for one representative TMA spot (donor no. 35377). Trichrome staining is shown for reference. Scale bar, 250 μm. b, CODEX antibody panel with 51 markers. c, Comparation of cells expressing both mesenchymal (Vimentin, αSMA) and epithelial (Keratin 8/18, E-cadherin, EpCAM) markers between Control and DN samples. d, Composition of immune cells in glomeruli-excluded tissue regions from DN kidneys (56 regions across 20 samples) and Control kidneys (6 regions across 2 samples).

**Extended Data Figure 2.**
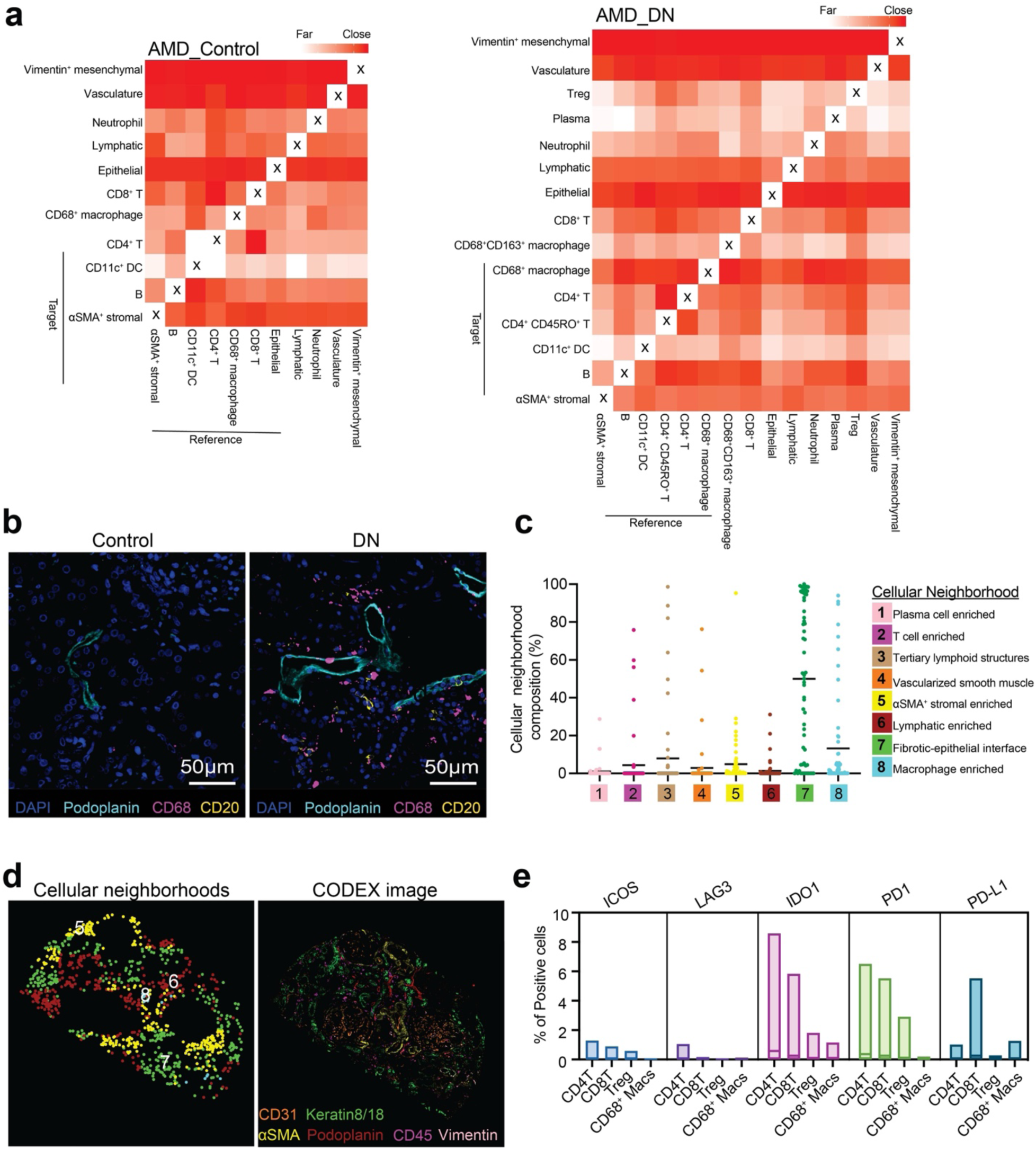
Cell colocalization and cell neighborhoods in CODEX dataset. a, Heatmap of average minimum distance (AMD) between reference cells and target cells in Control (left) and DN (right) samples. b, Representative regions of a Control and DN sample shown as four-color overlay images. Scale bar, 50 µm. c, Proportion of cells within distinct cell neighborhoods (CNs) relative to the total number of cells in each TMA core. Each dot represents the CN composition of an individual TMA core. d, Representative image of CELESTA cell type assignment (glomeruli-excluded) in 4 different colors to show 4 different CNs (left) and corresponding CODEX image (right) (donor no. 35381). e, Percentage of checkpoint-positive CD4^+^ T cells, CD8^+^ T cells, T regulatory (Treg) cells, and CD68^+^ macrophages in DN kidneys (56 regions across 20 samples). The line indicates the mean percentage.

**Extended Data Figure 3.**
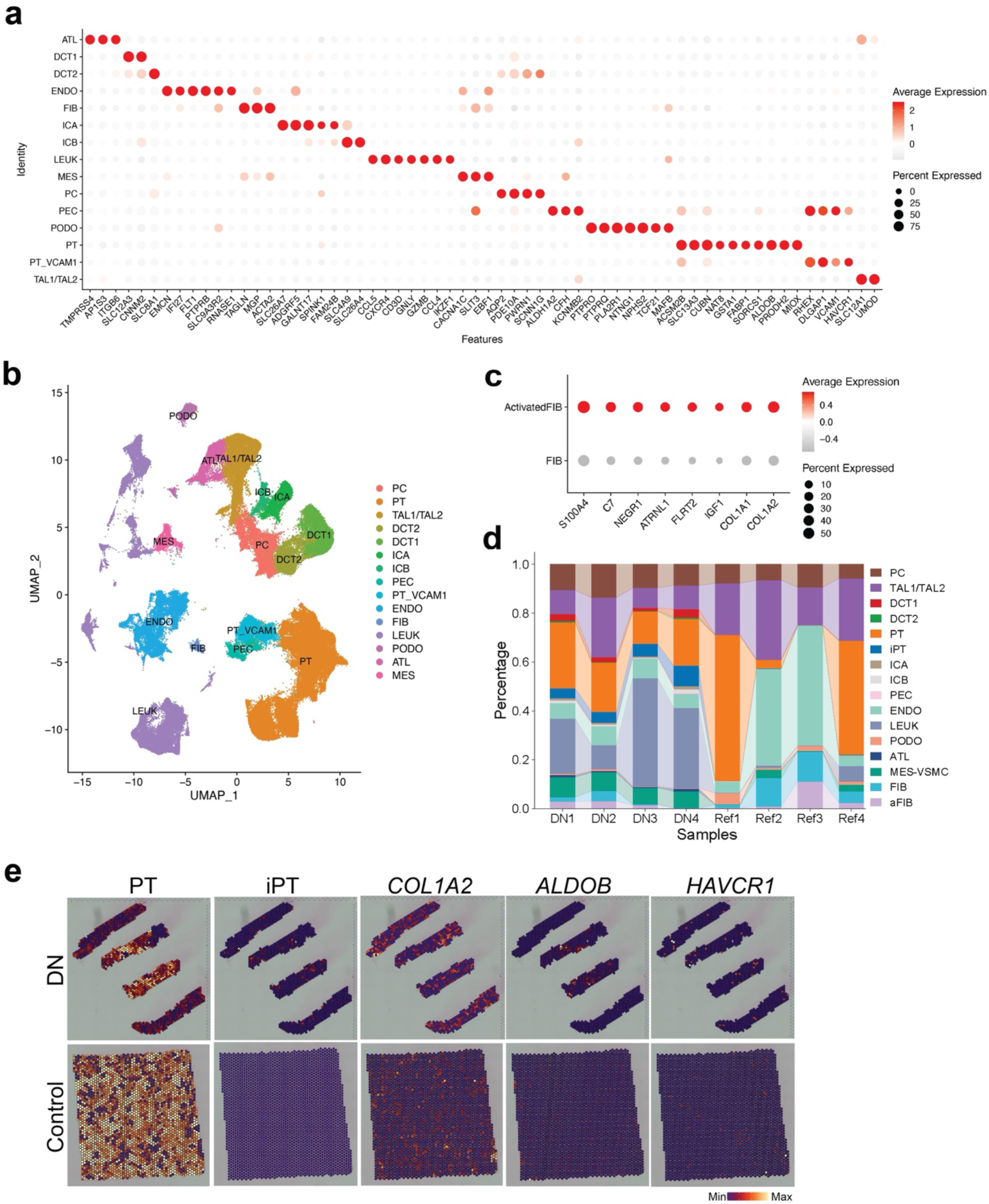
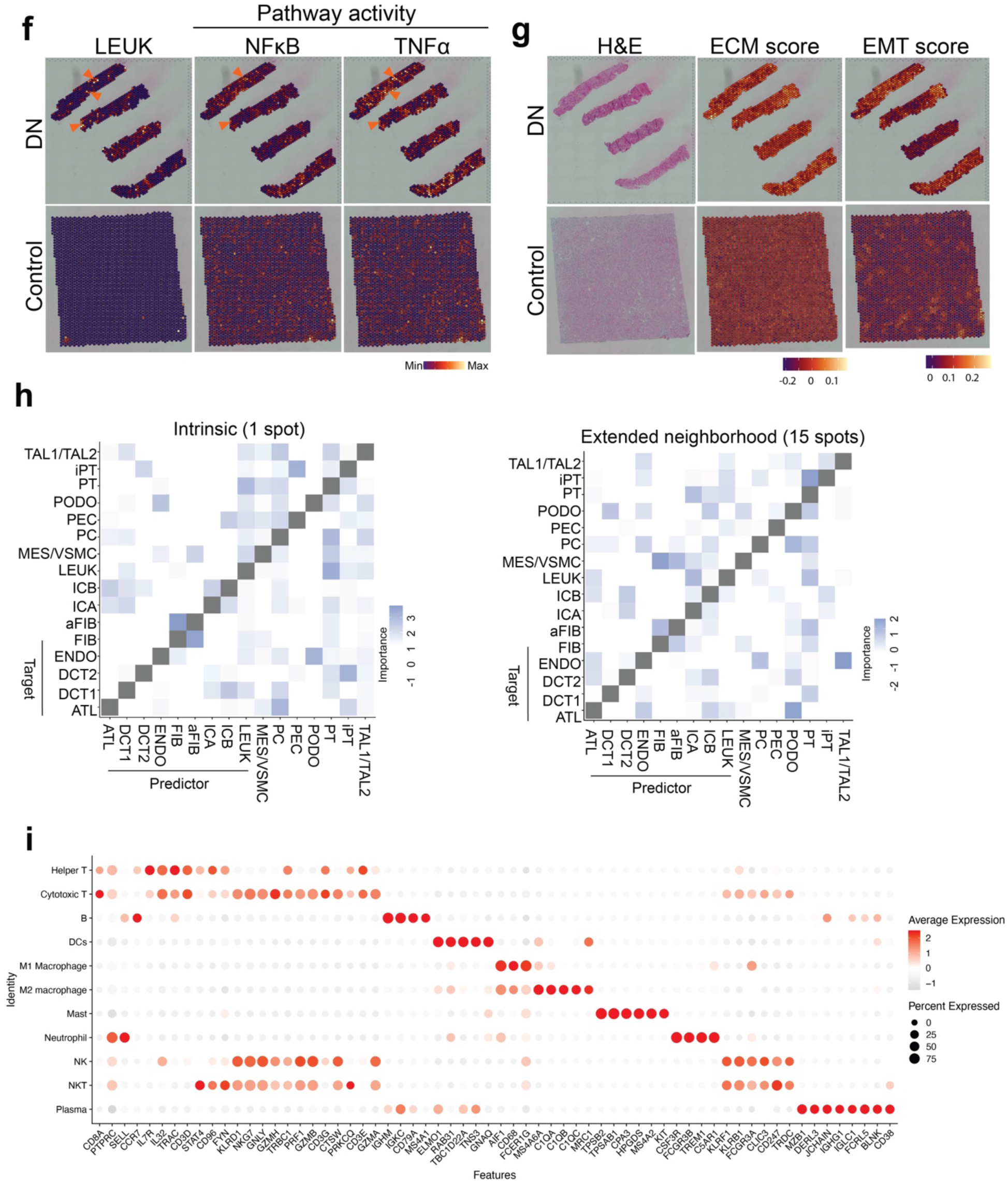
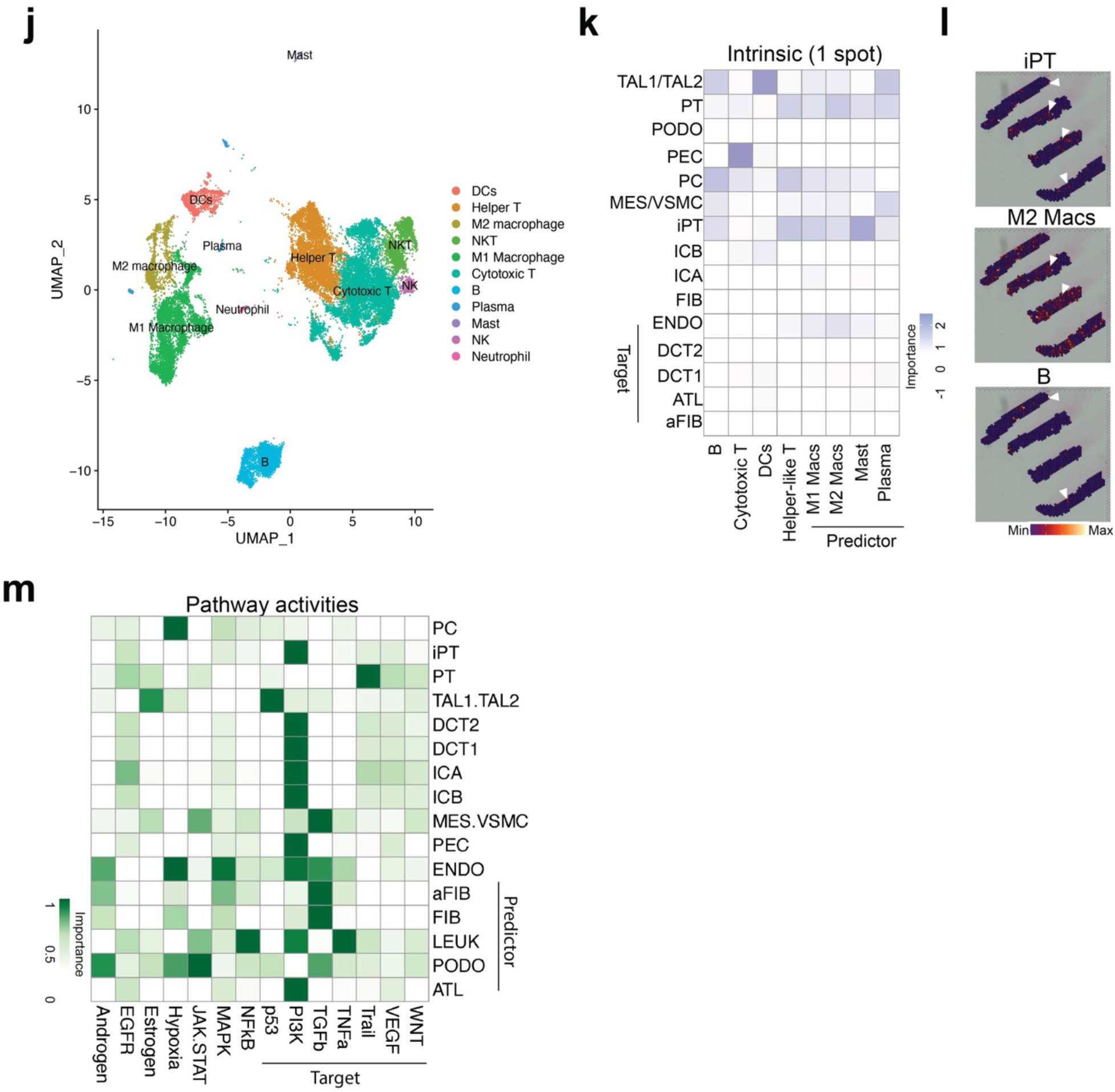
Comprehensive profiling of DN by integration of snRNAseq, scRNAseq and spatial transcriptomics datasets. a, Dot plots showing the major cell type marker gene average expression values. b, UMAP plot of the sc/snRNA-seq dataset showing major cell types. c, Dot plots showing the expression of marker genes of activated fibroblasts (aFIB). d, Bar plots showing cell-type proportions of sc/snRNA-seq dataset across all samples. Colors indicate different cell types. e, Visualization of PT, iPT cell types and gene expression of *COL1A2*, *ALDOB*, and *HAVCR1* in DN and Control tissues. Max, maximum; min, minimum. f, Visualization of LEUK cell type and their pathway activities in DN and Control tissues. Max, maximum; min, minimum. g, H&E staining and calculation of ECM and EMT scores in DN and Control tissues based on spatial mapping of ECM and EMT associated genes. h, Importance of the abundance of major cell types in the prediction of other cell types within a spot (intrinsic) and 15 spots (extended neighborhood). i, Dot plots showing the immune subpopulation marker gene average expression values. j, UMAP plot of the sc/snRNA-seq dataset showing immune subpopulations. k, Importance of the abundance of immune subtypes predict major cell types within a spot. l, Spatial mapping of iPT cells, M2 macrophages and B cells in a DN sample. Max, maximum; min, minimum. m, Importances of the abundance of major cell types in the prediction of PROGENy pathway activities within a spot.

**Extended Data Figure 4.**
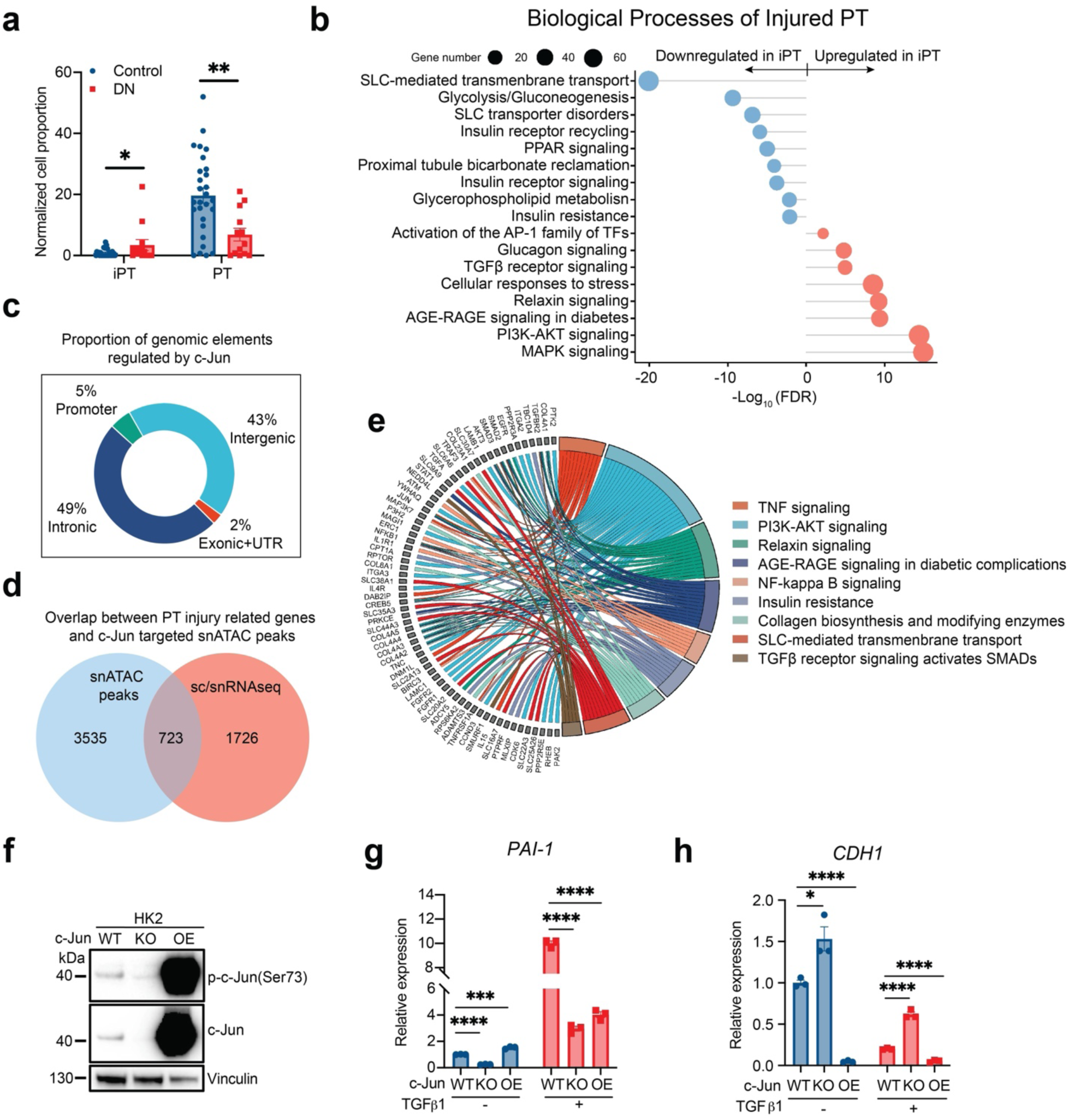
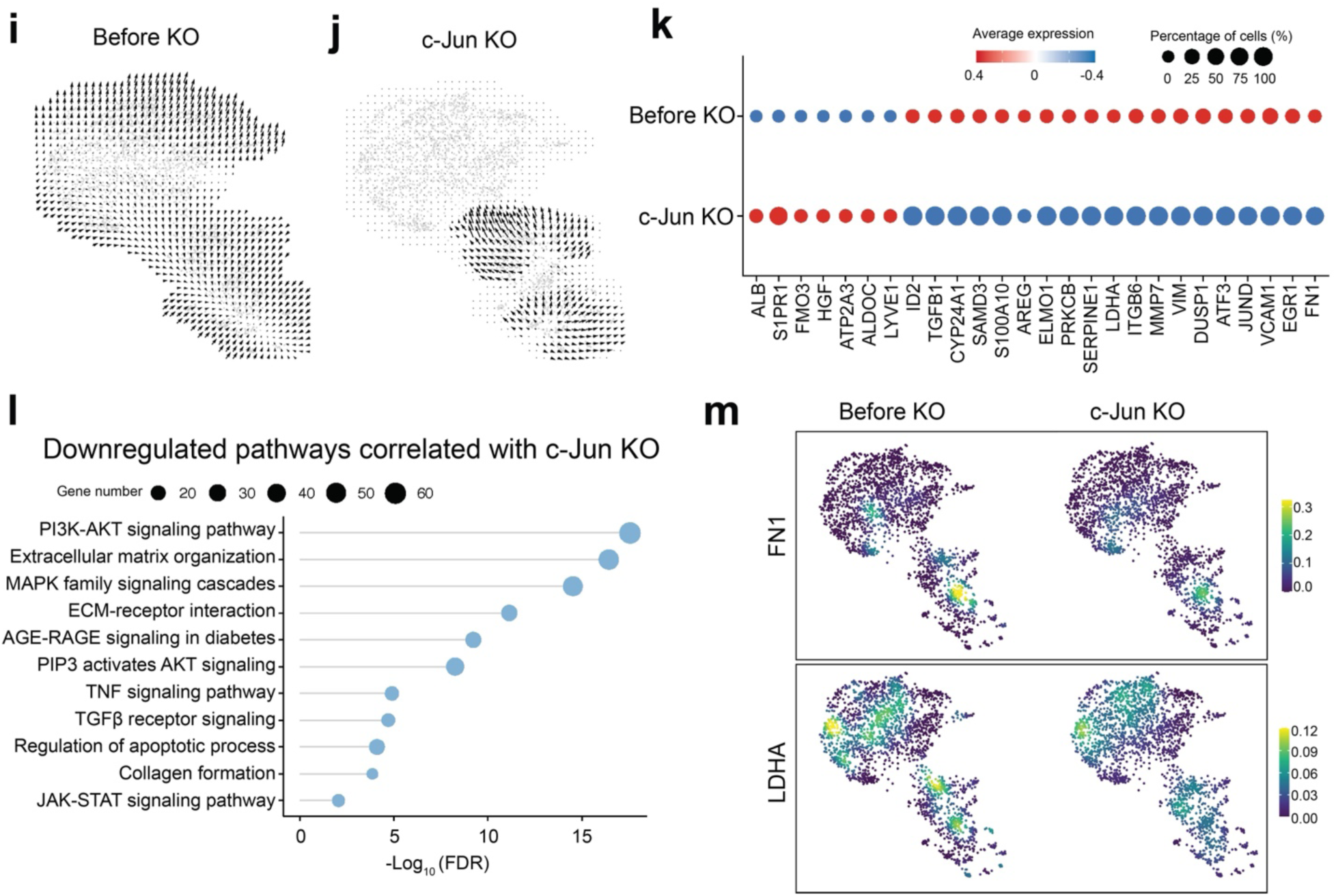
Characterization of c-Jun as a key regulator in sc/snRNAseq, snATACseq datasets, HK2 cells and *in-silico* knockout of *JUN* in PT cells. a, Cell proportion of iPT and PT cells in DN (n=13) and Control (n=28) samples. Each dot represents one sample. b, Pathway enrichment analysis for genes upregulated or downregulated in iPT cells compared to PT cells. c, Proportion of genomic elements regulated by c-Jun. d, Venn diagram illustrating the overlap between PT injury-associated genes identified by sc/snRNA-seq and genes associated with snATAC-seq peaks targeted by c-Jun. e, Pathway enrichment analysis for overlapped genes identified in d. f, Western blotting showing the expression of p-c-Jun, c-Jun expression in WT, c-Jun-KO and c-Jun-OE/HK2 cells. Vinculin represents as a loading control. g and h, mRNA expression of *PAI-1* and *CDH1* in WT, c-Jun-KO and c-Jun-OE/HK2 cells with or without TGFβ1treatment for 24 hours. Data are adjusted using unpaired two-tailed t-test. *P < 0.05, **P < 0.01, ***P < 0.001, ****P < 0.0001. i, RNA velocity combined with pseudotime analysis depicting the trajectory from PT to iPT states. j, *In-silico JUN* knockout disrupts the trajectory from PT to iPT. k, Expression of PT functional genes and the differential iPT markers before and after *JUN* knockout. l, Downregulated signaling pathways correlated with *JUN* knockout. m, Expression of representative iPT related genes before and after *JUN* knockout.

**Extended Data Figure 5.**
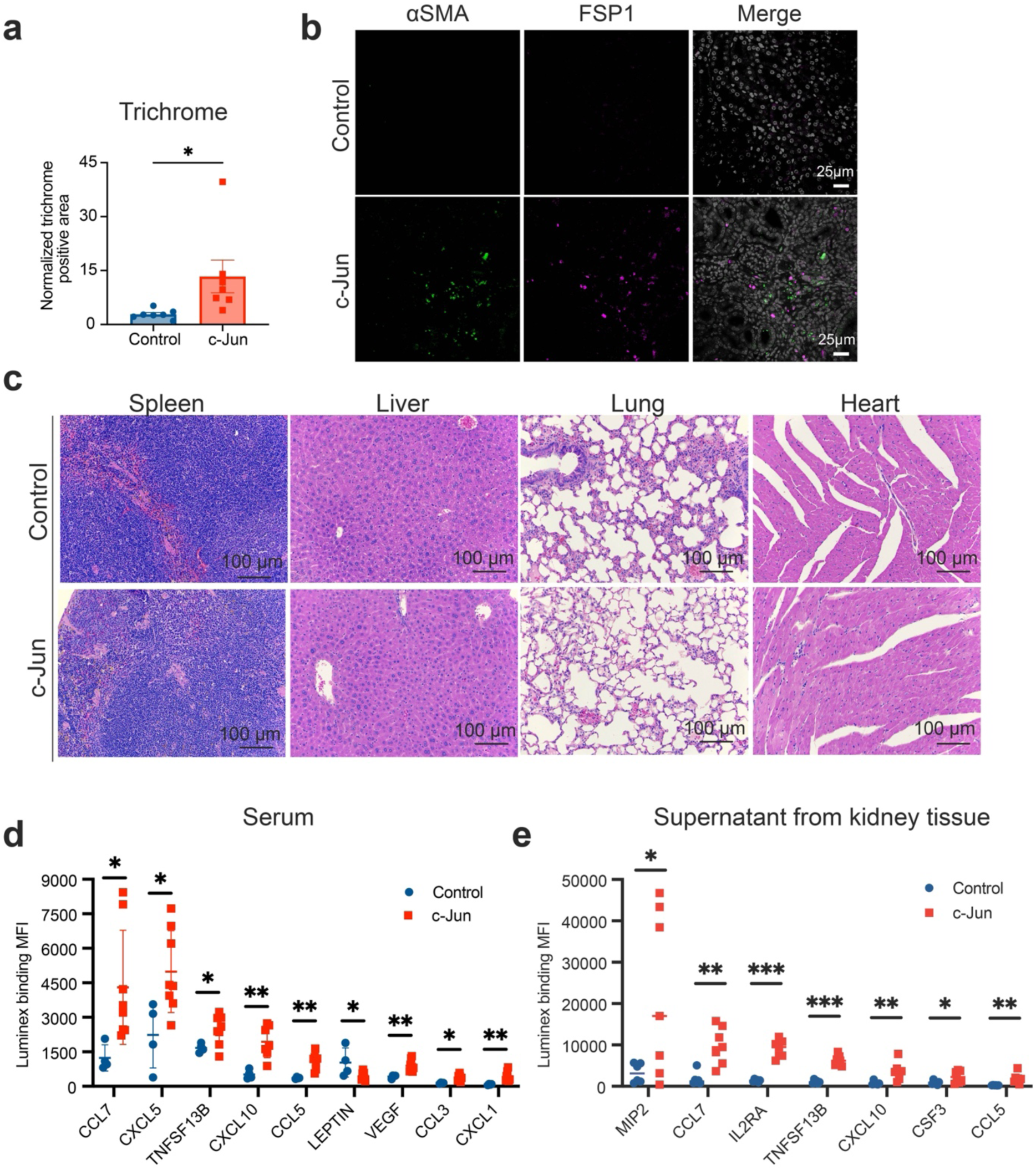
Characterization of c-Jun mice. a, Normalized trichrome positive area in control (n=7) and c-Jun (n=7) mice. b, Representative images of immunostaining for α-SMA and FSP1. Scale bar, 25 µm. c, Representative images of H&E staining of spleen, liver, lung and heart from indicated experimental groups. Scale bar, 100 µm. d, The secreted cytokines and chemokines in serum (Control, n=4; c-Jun, n=8) and e, the supernatant of kidney tissue homogenates (Control, n=6; c-Jun, n=7) of indicated experimental groups were quantified by Luminex assay. Data are represented as mean with S.D., adjusted using unpaired two-tailed t-test. *P < 0.05, **P < 0.01, ***P < 0.001, ****P < 0.0001.

**Extended Data Figure 6.**
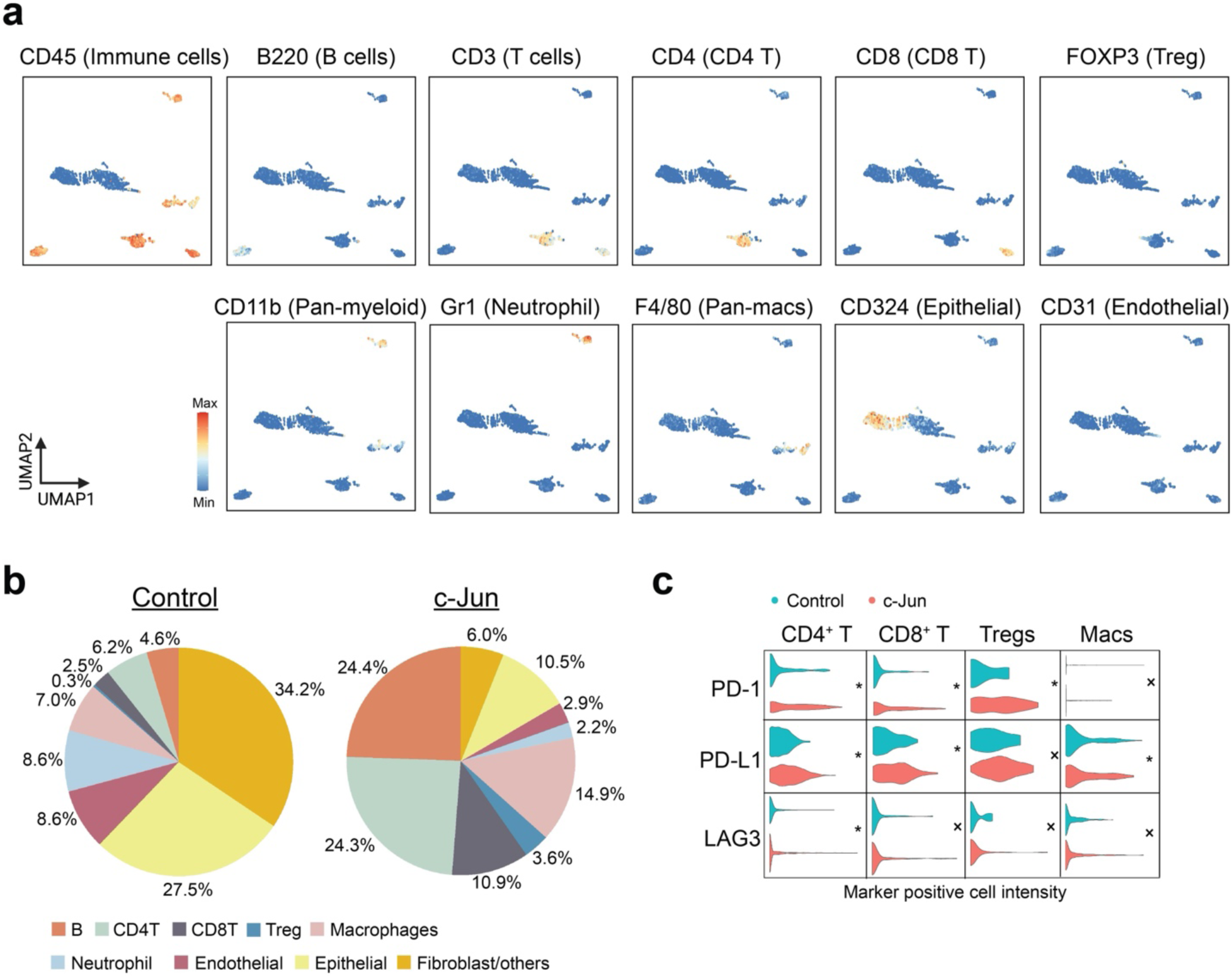
Immune phenotyping of c-Jun mice using CyTOF. a, UMAP plots displaying marker gene expression in the CyTOF dataset to differentiate cell types. b, Composition of cell types detected in control and c-Jun mice. c, Violin plots depict checkpoint-positive cell intensity levels across different immune subpopulation including CD4^+^ T cells, CD8^+^ T cells, Tregs, B cells and Macrophages in control and c-Jun mice. Data is normalized by ASINH ratio transformation to ensure comparability across varying intensity scales. Asterisks indicate statistically significant differences in marker-positive cell intensities between control and c-Jun mice, with significance levels denoted as follows: *P < 0.05, **P < 0.01, ***P < 0.001, and ****P < 0.0001, determined using Student’s t-test.

**Extended Data Figure 7.**
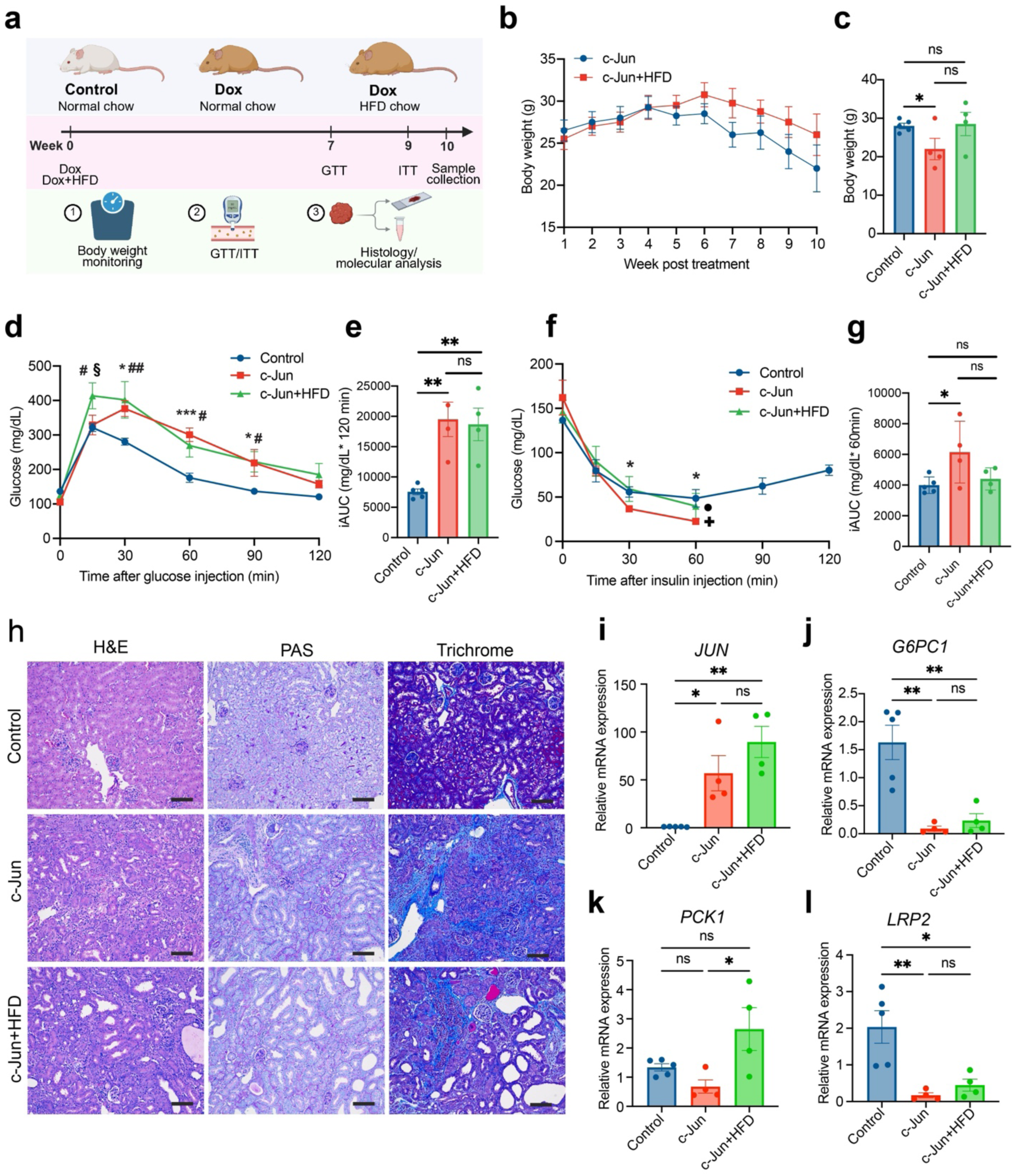
Tubular c-Jun activation perturbs systemic glucose metabolism and interacts with diet-induced metabolic stress. a, Timeline of the experimental design in male *c-Jun^tetO^ Pax8-rtTA* mice treated with Dox with or without a high-fat diet for 10 weeks. b, c, Body weight curves and endpoint body weight measurements showing weight changes among the groups.d, e, Glucose tolerance tests showing impaired glucose tolerance in c-Jun mice compared with controls, with no further worsening in c-Jun+HFD mice. “*” indicate P values for Control versus c-Jun; “#” indicate P values for Control versus c-Jun+HFD; “β” indicate P values for c-Jun versus c-Jun+HFD. Statistical comparisons were performed using ordinary 2way ANOVA. *(#, β)P < 0.05, **(##, ξξ)P < 0.01, ***(###, ξξξ)P < 0.001, ****(####, ξξξξ)P < 0.0001. f, g, Insulin tolerance tests showing an exaggerated hypoglycemic response in c-Jun mice that was partially blunted by HFD feeding. All control mice (n = 5) completed the ITT without intervention, whereas 3 of 4 c-Jun mice required glucose injection at the 60-min time point due to hypoglycemic symptoms (indicated by “+”). In the c-Jun+HFD group, 1 of 4 mice required glucose injection (indicated by “•”). “*” indicate P values for Control versus c-Jun. Statistical comparisons were performed using multiple t tests. *P < 0.05, **P < 0.01, ***P < 0.001, ****P < 0.0001. h, Representative H&E, PAS, and Trichrome staining showing comparable tubular injury and fibrosis in c-Jun and c-Jun+HFD kidneys. Scale bar, 50 µm. i–l, qPCR analysis of kidney tissue showing relative mRNA expression of *JUN*, *G6PC1*, *PCK1*, and *LRP2* in control (n=5), c-Jun (n=4), and c-Jun+HFD (n=4) mice. Data are presented as mean ± SEM. Statistical comparisons were performed using ordinary one-way ANOVA. *P < 0.05, **P < 0.01, ***P < 0.001, ****P < 0.0001.

**Extended Data Figure 8.**
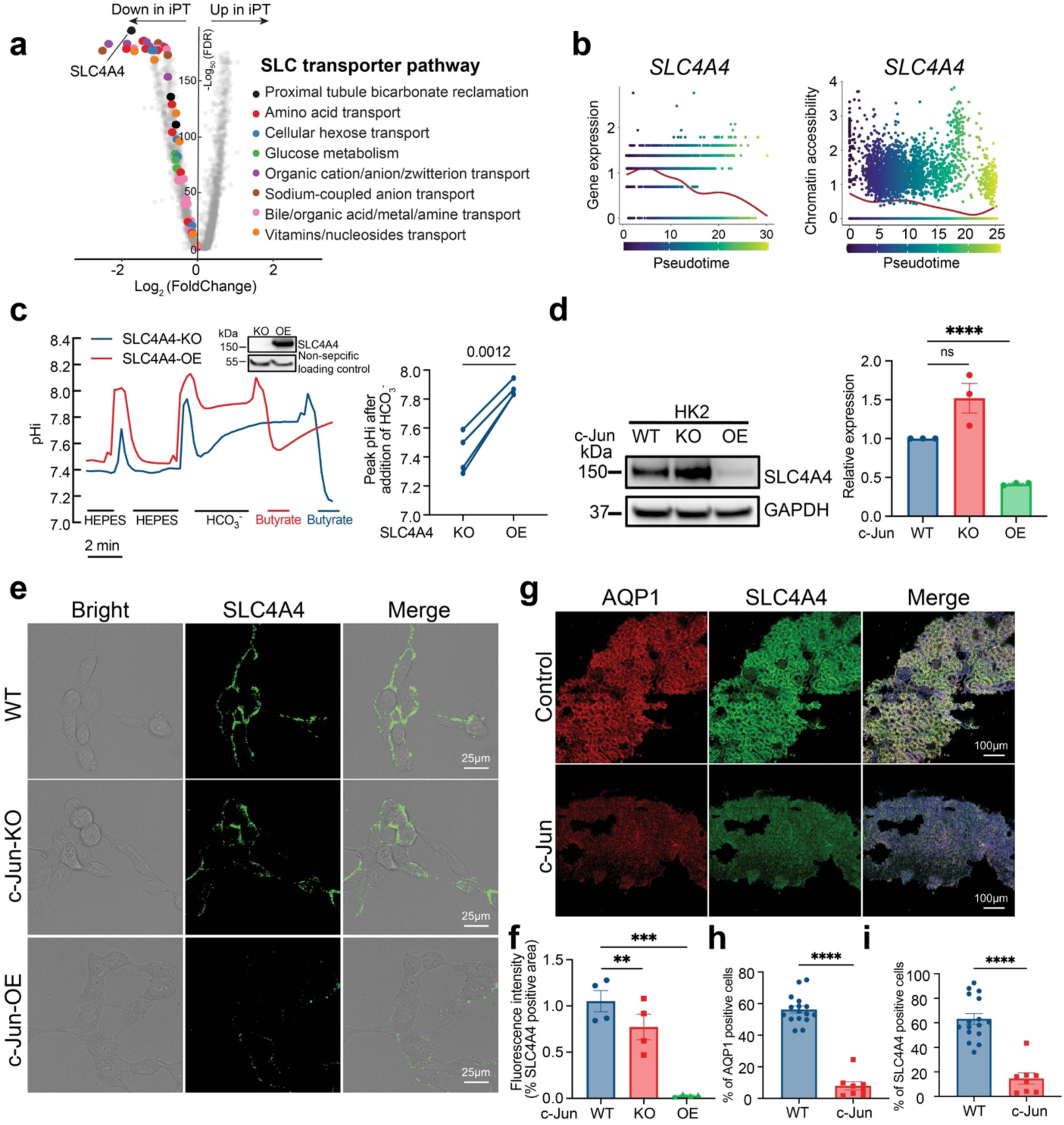
Downregulation of SLC4A4 in PT cells following c-Jun activation. a, Volcano plot showing differentially expressed genes associated with PT injury. Colored dots indicate downregulated SLC transporter genes, grouped by their associated transport pathways. b, Gene expression dynamics and chromatin accessibility dynamics of SLC4A4 along the pseudotime trajectory are shown. c, Representative trace of intracellular pH measurements in SLC4A4-KO and SLC4A4-OE cells in the presence of HEPES, HCO_3_^-^ and Butyrate (40 mM) buffered solution (left) and the summary of the changes of the intracellular pH at the peak level after addition of HCO_3_^-^ buffered solution (right). d, Western blotting showing the expression of SLC4A4 in WT, c-Jun-KO and c-Jun-OE/HK2 cells. The band intensities were analyzed by ImageLab (Bio-Rad), normalized by the expression in WT/HK2 cells. Each dot represents as an independent experiment. e, Immunostaining of the surface expression of SLC4A4 in live cells of WT, c-Jun-KO and c-Jun-OE/HK2 cells. Scale bar, 25 µm. f, Quantification of percentage of SLC4A4 positive area in four random fields are calculated. g, Immunostaining of AQP1 and SLC4A4 in Control and c-Jun mice. h, i, Quantification of the staining signals of AQP1 and SLC4A4. Percentages of AQP1 and SLC4A4-positive cells are shown. Each dot represents one mouse sample.

