## Supplementary Files for "A multi-omics map of diabetic nephropathy reveals c-Jun as a driver of tubular injury and metabolic stress"

**Supplementary Table 1. Demographic and clinical profiles of participants in CODEX assay**

| Donor ID | Organ | Age | Sex | Race/Ethnicity | Diabetes | Hypertension | Pathologic Classification (based on Tervaert et al, 2010) | % HbA1C | Serum albumin (g/dL) | EGFR (CKD-EPI formula without race) | Serum creatinine | Proteinuria, UPCR (mg/mg) | Proteinuria, urinalysis (mg/dL) | Proteinuria, dipstick | % Global glomerulo sclerosis | % IFTA | Extent of vascular disease (mild, moderate, severe arteriosclerosis) |
|---|---|---|---|---|---|---|---|---|---|---|---|---|---|---|---|---|---|
| DN1 | Kidney | 62 | F | N/A | YES | YES | IV | 6.6 | 3.4 | 9 | 4.87 | 14.72 | >300 | 3+ | 56 | 80 | Severe arteriosclerosis |
| DN2 | Kidney | 46 | M | Hispanic | YES | YES | IIb | 6.3 | 3.4 | 39 | 2.1 | 13 | >200 | 2+ | 47 | 30 | Severe arteriosclerosis |
| DN3 | Kidney | 67 | F | Hispanic | YES | YES | IIa | 6.6 | 4.2 | 13 | 3.6 | 1.15 | >100 | 2+ | 12 | 30 | Moderate arteriosclerosis |
| DN4 | Kidney | 45 | M | N/A | YES | YES | IV | 10.3 | 2.2 | 21 | 3.54 | 19.64 | >500 | 4+ | 89 | 90 | Severe arteriosclerosis |
| DN5 | Kidney | 67 | F | N/A | YES | YES | IV | 7.5 | 3.4 | 17 | 2.9 | 7.4 | >300 | 4+ | 78 | 80 | Moderate arteriosclerosis |
| DN6 | Kidney | 57 | M | Black/African America | YES | YES | IV | 7.7 | 2.6 | 19 | 3.65 | 9 | >100 | 2+ | 57 | 80 | Severe arteriosclerosis and hyalinosis |
| DN7 | Kidney | 39 | M | N/A | YES | YES | IV | 6.4 | 2.9 | 6 | 10.6 | N/A | >500 | 4+ | 78 | 80 | Severe arteriosclerosis and hyalinosis |
| DN8 | Kidney | 57 | M | Hispanic | YES | YES | III | 6.6 | 3.1 | 46 | 1.54 | 10.29 | >500 | 4+ | 37 | 30 | Severe arteriosclerosis |
| DN9 | Kidney | 81 | M | Chinese | YES | YES | IIa | 5.2 | 4.2 | 17 | 3.17 | 0.84 | >100 | 2+ | 44 | 35 | Moderate arteriosclerosis |
| DN10 | Kidney | 55 | F | Korean | YES | YES | III | 6.7 | 4.1 | 26 | 2.36 | 7 | >500 | 4+ | 30 | 60 | Severe arteriosclerosis |
| DN11 | Kidney | 38 | F | Black/African America | YES | YES | IV | 8.1 | 2.7 | 14 | 4.08 | 13.88 | >500 | 4+ | 71 | 80 | Moderate arteriosclerosis and extensive hyalinosis |
| DN12 | Kidney | 56 | M | Hispanic | YES | YES | III | 7.5 | 3.1 | 20 | 3.31 | 8.4 | >500 | 4+ | 12 | 80 | Arterial and arteriolar sclerosis and hyalinosis" |
| DN13 | Kidney | 55 | M | N/A | YES | YES | IV | N/A | N/A | 42 | 1.86 | 1.6 | N/A | N/A | 51 | 40 | Mild to moderate |
| DN14 | Kidney | 63 | F | N/A | YES | YES | IV | N/A | N/A | 18 | 2.9 | N/A | >300 | 3+ | 73 | 55 | Severe arteriolar hyalinosis |
| DN15 | Kidney | 49 | M | N/A | YES | YES | IV | 8.6 | 3 | 16 | 4.13 | N/A | >500 | 4+ | 63 | 90 | Arterial sclerosis and arteriolar hyalinosis |
| DN16 | Kidney | 59 | F | N/A | YES | YES | III | 14 | 2.6 | 58 | 1.05 | 6.79 | >500 | 3+ | 12 | 20 | Arteriolar hyalinosis and arterial sclerosis |
| DN17 | Kidney | 76 | F | N/A | YES | YES | III | 5.8 | 3.1 | 47 | 1.28 | 10.7 | >500 | 4+ | 11 | 10 | Arteriolar hyalinosis and arterial sclerosis |
| DN18 | Kidney | 35 | M | N/A | YES | YES | IV | 6.4 | 2.9 | 11 | 6.25 | 9.2 | >500 | 4+ | 77 | 90 | Mild arteriosclerosis |
| DN19 | Kidney | 56 | F | N/A | YES | YES | IV | 7.3 | 3.5 | 17 | 3.12 | N/A | N/A | N/A | 68 | 80 | Mild to moderate arteriosclerosis, severe arteriolar hyalinosis |
| DN20 | Kidney | 65 | F | N/A | YES | YES | III | 7.6 | 3.5 | 25 | 2.17 | 2.3 | >100 | 2+ | 25 | 35 | Mild arteriolosclerosis, focal arteriolar hyalinosis |
| Control1 | Lymph Node | | | | | | | | | | | | | | | | |
| Control2 | Kidney | | | | | | | | | | | | | | | | |
| Control3 | Tonsil | | | | | | | | | | | | | | | | |
| Control4 | Kidney | | | | | | | | | | | | | | | | |

**REFERENCE:** Tervaert, T. C. , Mooyaart, A. L. , Amann, K. , Cohen, A. H. , Cook, H. T. , Drachenberg, C. B. , Ferrario, F. , Fogo, A. B. , Haas, M. , de Heer, E. , Joh, K. , Noël, L. H. , Radhakrishnan, J. , Seshan, S. V. , Bajema, I. M. & Bruijn, J. A. (2010). Pathologic Classification of Diabetic Nephropathy. Journal of the American Society of Nephrology, 21 (4), 556-563. doi: 10.1681/ASN.2010010010.

**Supplementary Table 2. Demographic and clinical profiles of participants in Visium assay**

| Donor ID | Organ | Age | Sex | Race/Ethnicity | Diabetes | Hypertension | Pathologic Classification (based on Tervaert et al, 2010) | % HbA1C | Serum albumin (g/dL) | EGFR (CKD-EPI formula without race) | Serum creatinine | Proteinuria – UPCR (mg/mg) | Proteinuria – dipstick | % Global glomerulosclerosis | % IFTA | Extent of vascular disease (mild, moderate, severe arteriosclerosis) |
|---|---|---|---|---|---|---|---|---|---|---|---|---|---|---|---|---|
| DN1 | kidney | 65 | F | N/A | YES | YES | IV | 6.8 | 2 | 19 | 3.5 | 7.5 | 4+ | 50 | 60 | Severe |
| DN2 | kidney | 51 | M | Ukrainian | YES | YES | III | 9.8 | 1.5 | 39 | 1.95 | 9.73 | 3+ | 21 | 50 | Moderate to severe |
| DN3 | kidney | 53 | F | Pakistani | YES | YES | IV | 8.7 | 3.3 | 26 | 2.12 | >6 | 3+ | 60 | 40 | Severe |
| DN4 | kidney | 49 | M | Hispanic | YES | YES | III | 8.2 | 2.8 | 90 | 1.02 | 3.6 | 2+ | 9 | <5 | Arteriolar hyalinosis and arterial sclerosis |
| Control1 | kidney | 62 | F | N/A | NO | NO | | | | 85 | 0.85 | No proteinuria/negative | No proteinuria/negative | | | |
| Control2 | kidney | 74 | F | N/A | NO | NO | | | | 63 | 1.02 | No proteinuria/negative | No proteinuria/negative | | | |
| Control3 | kidney | 57 | M | White | NO | YES | | | | 58 | 1.52 | No proteinuria/negative | No proteinuria/negative | | | |
| Control4 | kidney | 56 | F | White | NO | YES | | | | 52 | 1.32 | No proteinuria/negative | No proteinuria/negative | | | |

**Supplementary Table 3. Antibody panel used in CyTOF assay**

| Mouse kidney | | | |
|---|---|---|---|
| Antigen | Mass | Element | Clone |
| BC1 | 102 | Pd | |
| BC2 | 104 | Pd | |
| BC3 | 105 | Pd | |
| BC4 | 106 | Pd | |
| BC5 | 108 | Pd | |
| BC6 | 110 | Pd | |
| CD45 | 89 | Y | 30-F11 |
| B220 | 111 | Cd | RA3-6B2 |
| Ki67 | 112 | Cd | B56 |
| Vimentin | 113 | In | D21H3 |
| αSMA | 115 | In | 1A4 |
| CD44 | 116 | Cd | IM7 |
| TER119 | 139 | La | Ter-119 |
| CD11b | 140 | Ce | M1/70 |
| CD31 | 141 | Pr | MEC13.3 |
| CD47 | 142 | Nd | miap301 |
| CD11c | 143 | Nd | HL3 |
| Gr1 | 144 | Nd | RB6-8C5 |
| CD4 | 145 | Nd | RM4-5 |
| Podoplanin | 146 | Nd | |
| CD3 | 147 | Sm | 17A2 |
| CD103 | 148 | Nd | 2.00E+07 |
| EPCAM | 149 | Sm | G8.8 |
| CD27 | 150 | Nd | LG.3A10 |
| ICOS | 151 | Eu | C398.4A |
| pAkt(S473) | 152 | Sm | D9E |
| CD274 (PD-L1) | 153 | Eu | 10F.9G2 |
| p-cJun | 154 | Sm | D47G9 |
| IDO | 155 | Gd | D5J4E |
| TIM-3 | 156 | Gd | RMT3-23 |
| CD19 | 157 | Gd | 6D5 |
| Foxp3 | 158 | Gd | FJK-16s |
| PD-1 | 159 | Tb | 29F.1A12 |
| CD206 | 160 | Gd | C068C2 |
| T-bet | 161 | Dy | O4-46 |
| CD163 | 162 | Dy | EPR19518 |
| CD80 | 163 | Dy | 16-10A1 |
| CD62L | 164 | Dy | MEL-14 |
| NK1.1 | 165 | Ho | PK136 |
| ARG1 | 166 | Er | poly-Rb |
| GATA3 | 167 | Er | TWAJ |
| CD8 | 168 | Er | 53-6.7 |
| CD63 | 169 | Tm | NVG-2 |
| E-cadherin (CD324) | 170 | Er | DECAM-1 |
| CD64 | 171 | Yb | X54-5/7.1 |
| CD86 | 172 | Yb | GL-1 |
| PanCK | 173 | Yb | AE1/AE3 |
| LAG-3 | 174 | Yb | C9B7W |
| CD273 | 175 | Lu | TY25 |
| F4/80 | 176 | Yb | T45-2342 |
| I-A/I-E (MHCII) | 209 | Bi | M5/114.15.2 |
| DNA | 191/193 | Ir | |
| Cisplatin Viability | 195 | Pt | |

| **Supplementary Table 4. List of RT-PCR primer sequences (human)** | | |
|---|---|---|
| Genes | Forward primer | Reverse primer |
| *FN1* | 5'-AGGCTTGAACCAACCTACGGATGA | 5'-GCCTAAGCACTGGCACAACAGTTT |
| *COL1A1* | 5'-CAATCACCTGCGTACAGAACGCC | 5'-CGGCAGGGCTCGGGTTTC |
| *TGF-β1* | 5'-TGGCGATACCTCAGCAACC | 5'-CTCGTGGATCCACTTCCAG |
| *PAI-1* | 5'-CACGAGTCTTTCAGACCAAG | 5'-AGGCAAATGTCTTCTCTTCC |
| *CDH1* | 5'-CGAGAGCTACACGTTCACGG | 5'-GGGTGTCGAGGGAAAAATAGG |
| *CGNL1* | 5'-CTGAGACTCGCAAGTGATGATAC | 5'-CCGAATACTGACACCGTAGGA |
| *SLC4A4* | 5'-TGATCGGGAGGCTTCTTCTCT | 5'-GGACCGAAGGTTGGATTTCTTG |
| *GAPDH* | 5'-ACCAGGGCTGCTTTTAACTCT | 5'-GGTGCCATGGAATTTGCC |
| *ACTB* | 5'-CATGTACGTTGCTATCCAGGC | 5'-CTCCTTAATGTCACGCACGAT |

| **List of RT-PCR primer sequences (mouse)** | | |
|---|---|---|
| Genes | Forward primer | Reverse primer |
| *JUN* | 5'-CCTTCTACGACGATGCCCTC | 5'-GGTTCAAGGTCATGCTCTGTTT |
| *G6PC1* | 5'-AGGTCGTGGCTGGAGTCTTGTC | 5'-GTAGCAGGTAGAATCCAAGCGC |
| *PCK1* | 5'-GGCGATGACATTGCCTGGATGA | 5'-TGTCTTCACTGAGGTGCCAGGA |
| *LRP2* | 5'-CCAATGGACTCACTCTGGACCT | 5'-GAATGGAAGGCAGTGCTGATGAC |
| *GAPDH* | 5'-ACAGTCCATGCCATCACTGC | 5'-GATCCACGACGGACACATTG |
| *TGF-β1* | 5'-ACTGATACGCCTGAGTGGCT | 5'-CCCTGTATTCCGTCTCCTTG |
| *FN1* | 5'-TGGTGGCCACTAAATACGAA | 5'-GGAGGGCTAACATTCTCCAG |
| *COL1A1* | 5'-GTGGTGACAAGGGTGAGACA | 5'-GAGAACCAGGAGAACCAGGA |

**Supplementary Table 5. The cell-type-signature matrix used in CELESTA based on the CODEX panel in the diabetic nephropathy study**

| Cell type | CD31 | CD34 | EPCAM | Karetin8/18 | aSMA | Podoplanin | Vimentin | CD45 | CD3e | CD20 | CD11c | CD163 | CD68 | CD66 | CD38 | CD8 | CD4 | CD45RO | FOXP3 | Round |
|---|---|---|---|---|---|---|---|---|---|---|---|---|---|---|---|---|---|---|---|---|
| Vasculature | 1 | 1 | 0 | 0 | 0 | 0 | NA | 0 | 0 | 0 | 0 | 0 | 0 | 0 | 0 | 0 | 0 | 0 | 0 | 1 |
| Epithelial | 0 | 0 | NA | 1 | 0 | 0 | NA | 0 | 0 | 0 | 0 | 0 | 0 | 0 | 0 | 0 | 0 | 0 | 0 | 1 |
| aSMA+ stromal | 0 | 0 | 0 | 0 | 1 | 0 | NA | 0 | 0 | 0 | 0 | 0 | 0 | 0 | 0 | 0 | 0 | 0 | 0 | 1 |
| Lymphatics | 0 | 0 | 0 | 0 | 0 | 1 | NA | 0 | 0 | 0 | 0 | 0 | 0 | 0 | 0 | 0 | 0 | 0 | 0 | 1 |
| Vimentin+ mesenchymal | 0 | 0 | NA | NA | NA | 0 | 1 | 0 | 0 | 0 | 0 | 0 | 0 | 0 | 0 | 0 | 0 | 0 | 0 | 1 |
| Immune | 0 | 0 | 0 | 0 | 0 | 0 | 0 | 1 | NA | NA | NA | NA | NA | NA | NA | NA | NA | NA | NA | 1 |
| CD3+ T cells | 0 | 0 | 0 | 0 | 0 | 0 | 0 | NA | 1 | 0 | 0 | 0 | 0 | 0 | NA | NA | NA | NA | NA | 2 |
| Neutrophils | 0 | 0 | 0 | 0 | 0 | 0 | 0 | NA | 0 | 0 | 0 | 0 | NA | 1 | NA | 0 | 0 | NA | 0 | 2 |
| B cells | 0 | 0 | 0 | 0 | 0 | 0 | 0 | NA | 0 | 1 | 0 | 0 | 0 | 0 | NA | 0 | 0 | NA | 0 | 2 |
| CD11c+ dendritic cells | 0 | 0 | 0 | 0 | 0 | 0 | 0 | NA | 0 | 0 | 1 | 0 | 0 | 0 | NA | 0 | 0 | NA | 0 | 2 |
| CD68+ macrophages | 0 | 0 | 0 | 0 | 0 | 0 | 0 | NA | 0 | 0 | NA | 0 | 1 | 0 | NA | 0 | 0 | 0 | 0 | 2 |
| CD68+CD163+ macrophages | 0 | 0 | 0 | 0 | 0 | 0 | 0 | NA | 0 | 0 | NA | 1 | 1 | 0 | NA | 0 | 0 | 0 | 0 | 2 |
| Plasma cells | 0 | 0 | 0 | 0 | 0 | 0 | 0 | NA | 0 | 0 | 0 | 0 | 0 | 0 | 1 | 0 | 0 | 0 | 0 | 2 |
| CD8+ T cells | 0 | 0 | 0 | 0 | 0 | 0 | 0 | NA | NA | 0 | 0 | 0 | 0 | 0 | NA | 1 | 0 | NA | NA | 3 |
| CD4+ T cells | 0 | 0 | 0 | 0 | 0 | 0 | 0 | NA | NA | 0 | 0 | 0 | 0 | 0 | NA | 0 | 1 | NA | NA | 3 |
| CD4+ CD45RO+ T cells | 0 | 0 | 0 | 0 | 0 | 0 | 0 | NA | NA | 0 | 0 | 0 | 0 | 0 | NA | 0 | NA | 1 | NA | 4 |
| Tregs | 0 | 0 | 0 | 0 | 0 | 0 | 0 | NA | NA | 0 | 0 | 0 | 0 | 0 | NA | 0 | NA | NA | 1 | 4 |
